# Diverse mechanisms control amino acid-dependent environmental alkalization by *Candida albicans*

**DOI:** 10.1101/2023.08.04.551922

**Authors:** Fitz Gerald S. Silao, Valerie Diane Valeriano, Erika Uddström, Emilie Falconer, Per O. Ljungdahl

## Abstract

*Candida albicans* has the remarkable capacity to neutralize acidic growth environments by releasing ammonia derived from the catabolism of amino acids. The molecular components and mechanisms controlling this capacity remain poorly understood. Here, we present an integrative model with the cytosolic NAD^+^-dependent glutamate dehydrogenase (Gdh2) as the principal component. We show that the alkalization defect of a strain lacking the SPS-sensor regulated transcription factor *STP2* is due to the inability to fully derepress *GDH2* and the two proline catabolic enzymes, *PUT1* and *PUT2*. Notably, the Stp2-dependent regulation of *PUT1* and *PUT2* occurs independent of Put3, the proline-dependent activator. Accordingly, a *stp2-/– put3-/-* strain is unable to derepress the expression of these enzymes resulting in a severe alkalization defect that nearly phenocopies the abrogated alkalization of a *gdh2-/-* strain. In wildtype cells, alkalization is tightly dependent on mitochondrial activity and occurs as long as conditions permit respiratory growth. As alkalization proceeds, Gdh2 levels decrease and glutamate is transiently extruded from cells. Together these two processes constitute a rudimentary regulatory system enabling cells to prevent the rapid intracellular build-up of ammonia. Similar to *C. albicans*, Gdh2-dependent alkalization is dispensable for *C. glabrata* and *C. auris* virulence as assessed using a whole-blood infection model. Intriguingly, fungal-dependent alkalization does not influence the growth or proliferation of *Lactobacillus crispatus*, a potent antagonist of *C. albicans* that normally resides in the acidic vaginal microenvironment. Our data suggest that it is time to reconsider the idea that pH modulation driven by pathogenic fungi plays a crucial role in shaping the architecture and dynamics of (poly)microbial communities. Other factors are likely to be more critical in contributing to dysbiosis and that favor virulent growth.

## Introduction

*Candida albicans* is the primary cause of human mycoses presenting a spectrum of pathologies ranging from superficial lesions to life-threatening invasive infections. Despite its ill repute, *C. albicans* is normally a harmless commensal, thriving as a benign member of human microbiota in healthy individuals. Amino acids are among the most abundant nutrients in human hosts capable of supporting growth of this fungus. Yet, the utilization of amino acids in excess of the amount necessary for growth and to support basic cellular functions must be controlled by *C. albicans* due to the risk of accumulating ammonia (NH_3_), a weak base that becomes toxic when in excess [1]. The release of ammonia (NH_3_) into the extracellular space alkalinizes the growth environment as it converts to ammonium (NH_4_ ^+^) [2]. The interest in studying this process in *C. albicans* has been linked to findings that alkaline pH induces filamentous growth in vitro; morphological switching is a known virulence characteristic. Extracellular alkalization via ammonia release is not exclusive to *C. albican*s, other members of the pathogenic *Candida* species complex also are capable of alkalizing their growth environments [2]. *C. albicans*, *C. auris,* and *C. glabrata* are among the top priority fungal pathogens listed by the World Health Organization (WHO), with *C. auris* and *C. albicans* placed under the “critical priority” group [3, 4].

We recently showed that the NAD^+^-dependent glutamate dehydrogenase (Gdh2; EC:1.4.1.2), which catalyzes the deamination of glutamate to α-ketoglutarate [5–7], is the key enzyme responsible for ammonia generation during growth on amino acids [6]. Deletion of *GDH2* completely abrogates the ability of strains to alkalinize all amino acid-based media examined, in dense cultures (OD ≍ 5) and even during extended periods of incubation [6]. This defect surpassed the prominent alkalization deficiency observed in a strain lacking *STP2* [6], which has been extensively referenced as an alkalization-deficient strain [2, 8–13]. *STP2* encodes a transcription factor required to derepress the expression of several amino acid permease genes and a subset of oligopeptide transporters [10, 14]. Stp2 is produced as a latent precursor that is proteolytically activated by the concerted action of the SPS (<u>S</u>sy1-<u>P</u>tr3-<u>S</u>sy5) sensor in response to the presence of extracellular amino acids.

The processed form of Stp2, lacking an N-terminal cytoplasmic retention domain, efficiently translocates to the nucleus where it binds to promoters of target genes [5, 10, 14]. In addition to alkalization deficiency, the *stp2* strain exhibits reduced virulence in a murine systemic infection model and in mouse bone marrow-derived macrophages [8, 15, 16], which rationally led to the conclusion that the inability to alkalinize and reduced virulence are linked. However, this notion is inconsistent with our data, as the inactivation of *GDH2* was found not to affect the virulence of *C. albicans* in a murine systemic infection model and did not prevent morphological switching of cells phagocytized by macrophages [6]. Thus, Stp2-dependent virulence must be linked to other alkalization-independent processes.

Despite these new insights, a clear mechanistic understanding is lacking of how environmental alkalization is achieved and controlled when *C. albicans* utilizes amino acids for growth. For instance, in addition to the known effectors of alkalization (Gdh2, Stp2), it is established that media alkalization depends on cell density. In fact, abrogation or a significant delay in alkalization in dense liquid cultures (e.g., OD ≥ 2) grown over an extended period is an “acid test” for alkalization requirement, and only a handful of mutant strains are known to pass this test. It is also established that mitochondrial function is key to effect alkalization, which only occurs when cells are grown under respiratory conditions, i.e., medium with low (<0.2%) glucose or alternative carbon sources, such as glycerol and/or lactate [2, 6]. Interestingly, the <u>P</u>roline <u>UT</u>ilization (PUT) pathway, a major pathway for the generation of glutamate, is also repressed by high glucose. The PUT pathway is comprised of Put1 (proline dehydrogenase; EC 1.5.5.2) and Put2 (Δ1-pyrroline-5-carboxylate (P5C) dehydrogenase; EC 1.2.1.88) that are localized exclusively in the mitochondria [6, 17]. Strains lacking *PUT1* and *PUT2* exhibit significantly reduced capacities for alkalinizing the growth media, indicating the presence of other pathways to generate glutamate [5, 6]. Interestingly, in contrast to mammalian cells, where the NAD^+^-dependent glutamate dehydrogenase (GDH) is localized to the mitochondria and the α-ketoglutarate formed from glutamate feeds directly into the TCA cycle, Gdh2 in *C. albicans* is a cytoplasmic enzyme [6, 17]. This difference raises the question as to how *C. albicans* cells coordinate reactions occurring in spatially exclusive sites to properly control the effects of environmental alkalization.

Based on findings reported here, we have developed an integrative model that accounts for how *C. albicans* generates ammonia and initiates neutralization of an acidic environment. We posited that Gdh2 is the critical component, which enabled us evaluate the contributions and connections of other known factors linked to amino acid uptake and mitochondrial function. Consistent with our finding in *C. albicans*, we show that although *GDH2* is essential for amino acid dependent-alkalization in other pathogenic *Candida* species, such as *C. glabrata* and *C. auris*, it remains dispensable for virulence as assessed by survival in whole blood culture. Furthermore, by examining the proliferation of *Lactobacillus crispatus*, a potent antagonist of *C. albicans* that normally thrives in the acidic vaginal microenvironment, we found that environmental alkalization does not directly limit the growth of this competing microorganism.

## Results

### Gdh2 level is sensitive to extracellular pH

We have reported that Gdh2 levels decrease as the pH of the growth medium increases towards neutrality [6]. To directly examine the effect of pH on Gdh2 stability, we followed the Gdh2 levels 1 h after cultures were transferred to buffered media with defined pH (pH 4 – 8) and spiked with cycloheximide (CHX) to arrest translation (Fig. 1A). The strain (CFG433) was pre-grown in YPD and shifted to alkalization media, a synthetic medium containing casamino acids (CAA) as sole carbon/nitrogen/energy source (YNB+CAA; pH = 4, unbuffered), for 2 h to induce the expression of Gdh2. CAA, derived from the controlled acid hydrolysis of casein, contains free amino acids and oligopeptides at a ratio of 82% to 18%, respectively [18]. Relative to T = 0 h, there was no significant change in Gdh2 levels at pH = 4, indicating that Gdh2 is relatively stable at acidic pH (Fig. 1B). Interestingly, a significant drop in the levels of Gdh2 was observed at pH = 5. Consistently, the levels of Gdh2 in the unbuffered culture, where the pH ∼5 (see wells in Fig. 1A), were also significantly reduced after 1 h. Since the strain also co-expresses Put1-RFP and Put2-HA, we analyzed the levels of these enzymes in the same lysates to gather more information related to the interaction of these spatially separated enzymes during alkalization. Both Put1 and Put2 were stable under all the pH conditions tested (Fig. 1B), suggesting that glutamate generation in the mitochondria from proline catabolism remains active and that the reduced levels of Gdh2 in the cytosol may provide a mechanism of controlling the intracellular build-up of glutamate-derived ammonia as the external pH increases.

**Fig. 1.**
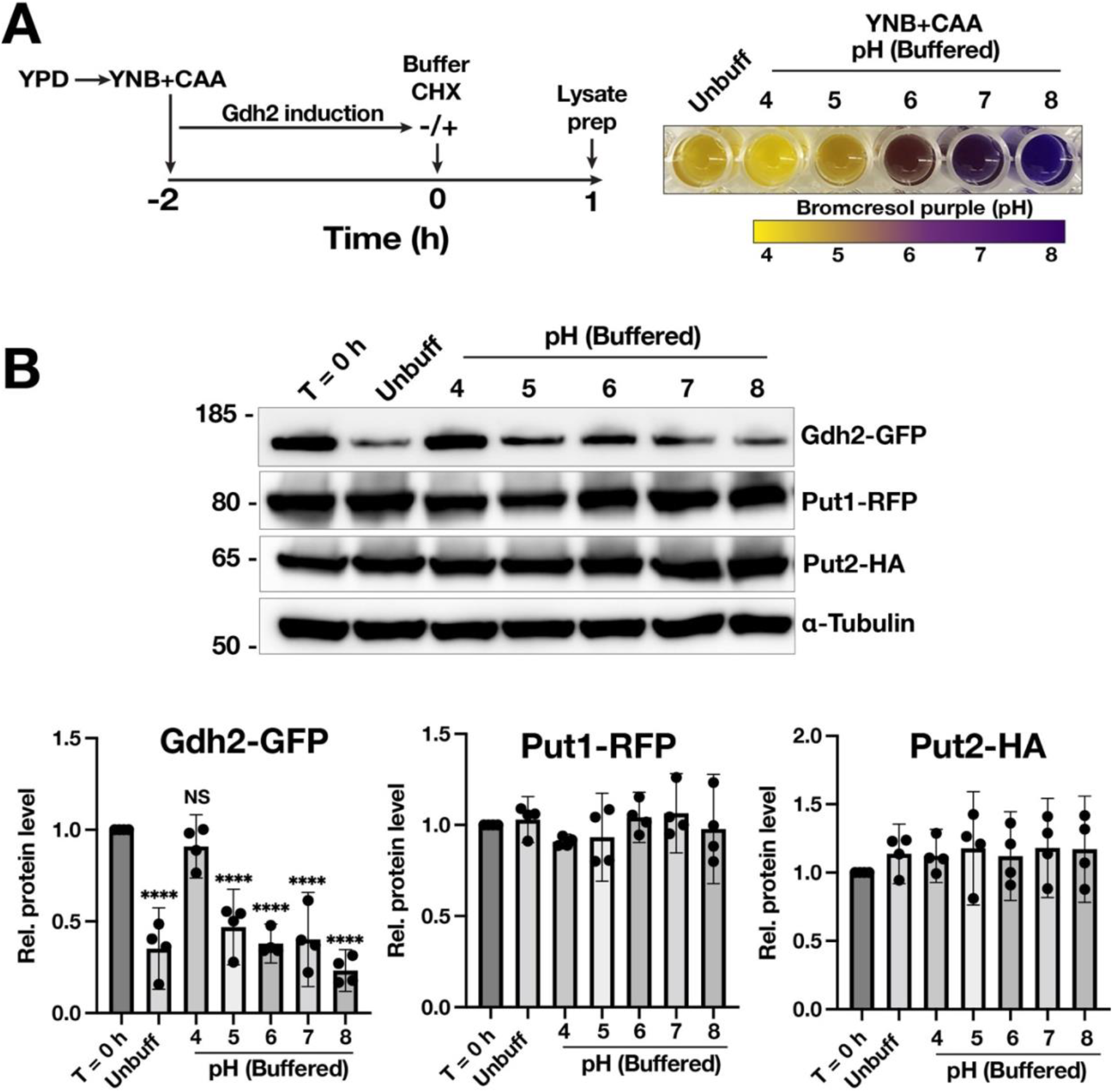
Gdh2 level is sensitive to extracellular pH. A) Scheme for the alkalization experiment. Gdh2 was induced for 2 h in YNB+CAA with bromocresol purple (BCP) as pH indicator and then spiked with concentrated buffer to 50 mM final concentration and cycloheximide (CHX) to arrest translation. Cultures were incubated for another 1 h followed by whole cell lysis and immunoblotting. (Right panel) Aliquots of spiked cultures after 1 h were placed in microplate wells to show the change in color corresponding to the indicated pH. **B)** Immunoblot and quantification of Gdh2-GFP, Put1-RFP, and Put2-HA levels in cells grown at different culture pH. The signals from the target proteins were first normalized to α-tubulin and then compared relative to T = 0h (set to 1). Data presented are from 4 biological replicates (mean with 95% CI; analyzed by one way ANOVA followed by Dunnett’s posthoc test, \*\*\*\**p* <0.0001).

### High glucose and mitochondrial activity regulate glutamate availability for Gdh2

Previous studies have shown that high levels of glucose (2%) in a CAA-based medium inhibit extracellular alkalization [2]. However, Gdh2 is weakly expressed YNB+CAA with 2% glucose [6], although at levels significantly lower than in a medium lacking glucose (Fig. 2A). Consequently, cells growing in the presence of high glucose, should retain a limited capacity to produce ammonia and to alkalinize the medium. To reconcile these findings, we posited that the substrate glutamate becomes limiting due to glucose repression of processes that generate it. We have previously shown that the components of the proline utilization (PUT) pathway, Put1 and Put2, that generate glutamate in the mitochondria and are repressed by high levels of glucose [5, 6]. Immunoblotting analysis showed that, like Gdh2, Put2 is also significantly reduced in medium containing high glucose (Fig. 2A). Consistently, the level of intracellular glutamate is significantly lower in cells grown in alkalization medium containing 2% glucose compared to medium lacking it (Fig. 2B), which clearly suggests that in addition to repressing the levels of Gdh2, high glucose also limits the production of its substrate, and hence ammonia production.

**Fig. 2.**
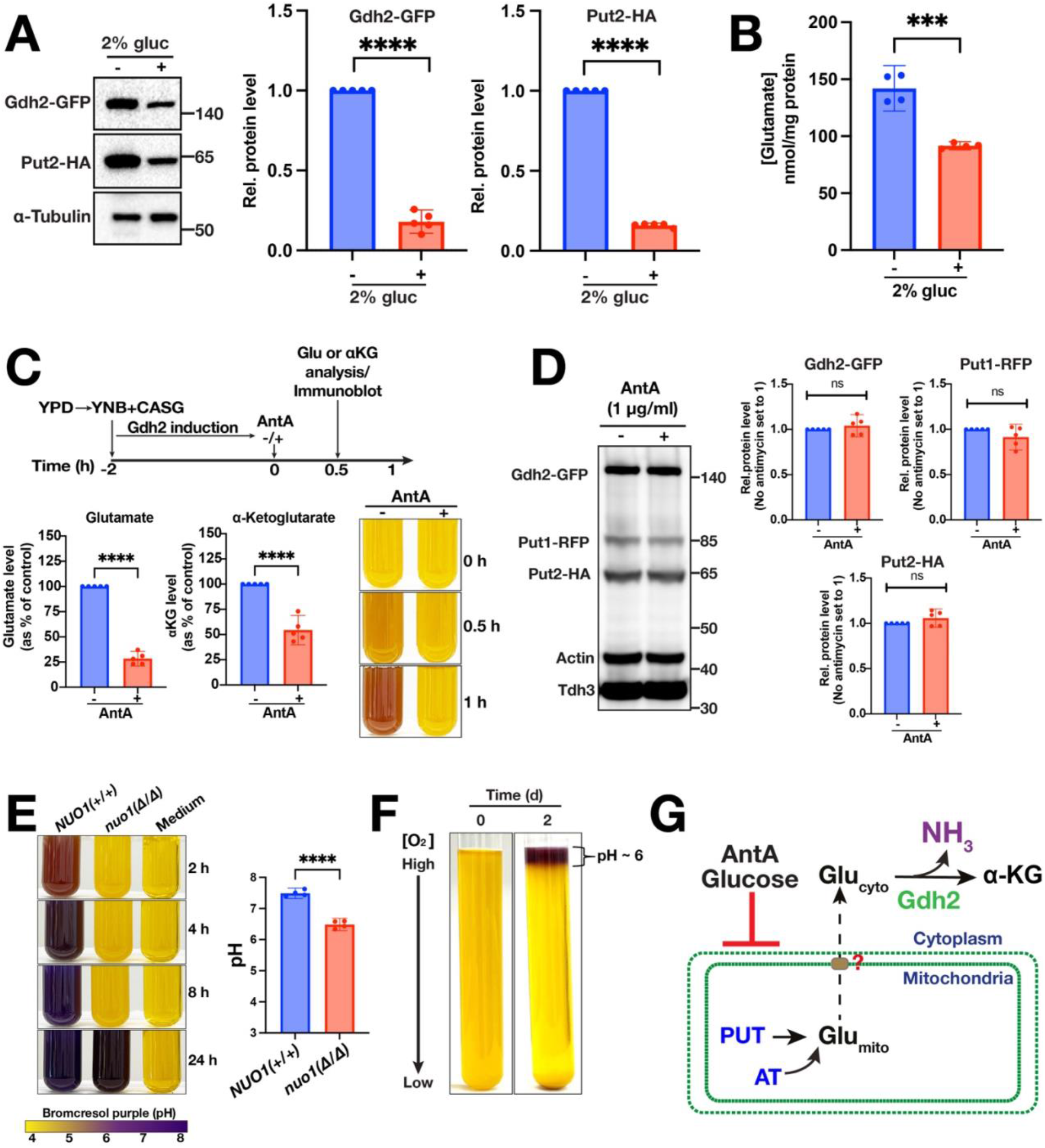
High glucose and mitochondrial activity regulate glutamate availability for Gdh2. A) Immunoblot analysis of Gdh2-GFP and Put2-HA levels in lysates prepared from cells (CFG404) grown in YNB+CAA without or with 2% glucose. **B)** Intracellular glutamate levels in lysates prepared from (A). Data shown were obtained from 4-5 biological replicates (mean with 95% CI; \*\*\**p* <0.0001 by student *t*-test). **C**) Antimycin A (AntA) arrests alkalization. Cells were grown for 2 h in YNB+CAA medium to induce Gdh2 expression and then spiked with AntA or vehicle control (ethanol, –). Tubes were photographed at the indicated timepoints (Right) or sampled after 30 min to determine the levels of cytosolic glutamate and α-ketoglutarate (Left); results are presented as percentage (%) of vehicle control derived from 5 biological replicates (mean with 95% CI; \*\*\*\**p* <0.0001 by student *t*-test). **D**) Immunoblot analysis of cells grown and sampled as in (**C**). Enzymes levels are not altered after exposing the cells to AntA. Enzymes were simultaneously detected in the same lysate and quantified using Tdh3 as loading control. Data is presented as mean with 95% CI (n = 5). **E**) Strain with mitochondrial dysfunction exhibits striking alkalization defect. (Left) Wildtype *NUO1+/+* (SN250) and *nuo1*Δ/Δ (CB342) strains were grown in YNB+CASG at OD ≍ 5 and photographed at the indicated timepoints. (Right) pH values of cultures after 24 h (mean with 95% CI; \*\*\*\**p* <0.0001 by student *t*-test). **F)** CFG441 cells were inoculated into 20 ml of YNB+CAA at OD ≍ 5 and then kept upright at room temperature for 2 days. Only the top layer with the highest oxygen concentration resulted to alkalization. **G)** Schematic diagram of AntA or high glucose inhibits alkalization. AntA or high glucose inhibits mitochondrial function and prevent the export of Glu_mito_ generated by either proline catabolism (PUT) or aminotransferases (AT) to the cytosol.

In addition to limiting the levels of glutamate, high glucose pleiotropically represses mitochondrial activity, which can affect the movement of substrates between the cytosol and the mitochondria [19]. Consistently, in previous work, we showed that alkalization was arrested in glucose-devoid cultures (OD ≍ 5) treated with the potent mitochondrial complex III inhibitor antimycin A (AntA) [6], supporting the notion that mitochondrial activity is essential to alkalization. We posited that the bulk of the glutamate in the cytosol originates from the mitochondria and that acute mitochondrial inhibition during alkalization reduces the cytoplasmic glutamate pool. To test this, we analyzed the level of glutamate in the cytosol of the reporter strain (CFG441) grown in YNB+CASG (i.e., YNB+CAA medium supplemented with 38 mM ammonium sulfate and 1% glycerol) 30 min after it was treated with AntA (1 μg/ml) (Fig. 2C, top panel). The cytosol fraction was extracted using the procedure reported by Ohsumi et al. [20] using dilute CuCl_2_ (0.2 mM) to gently permeabilize the plasma membrane. We observed a significant ∼3-fold reduction in the level of cytosolic glutamate following treatment by AntA (Fig. 2C). As expected, alkalization was arrested by AntA as inferred from the color of the pH indicator, while that of the carrier control (ethanol) showed continued alkalization (Fig. 2C). Consistent to glutamate being limiting, the level of α-ketoglutarate was significantly reduced (Fig. 2C) despite no alteration in the levels of Gdh2, Put1, and Put2 (Fig. 2D). We deliberately limited the analysis of expressed enzyme levels to 30 min after the addition of AntA, as the pH of the unbuffered control is expected to increase and independently reduce the Gdh2 levels (Fig. 1B).

To further test the role of mitochondria in alkalization, we examined the alkalization capacity of a strain lacking *NUO1*, which encodes a subunit of mitochondrial NADH:ubiquinone oxidoreductase (complex I). The *nuo1*Δ/Δ manifests the classical mitochondrial-deficient phenotypes, including reduced respiration (oxygen consumption), low mitochondrial membrane potential, and extremely poor growth on non-fermentable carbon sources [21, 22]. Considering this, we grew the cultures in YNB+CASG at starting OD ≍ 5. The *nuo1*Δ/Δ strain was unable to alkalinize the medium after 8 h, but eventually the media became alkaline upon prolonged (24 h) incubation (Fig. 2E)[6]. Finally, a simple classical experiment based on allowing dense cultures to grow static at room temperature, alkalization was only observed at the top layer that is exposed to the atmosphere, which clearly links alkalization to respiratory growth (Fig. 2F). Taken together, these results support the scheme shown in Fig. 2G where the mitochondria actively contributes to the activity of Gdh2 by supplying the cytosol with mitochondrial-derived glutamate produced via proline catabolism (PUT) and/or some other mitochondrial-localized aminotransferases (AT) [19].

### Glutamate excretion as a mechanism to regulate the intracellular glutamate pool size

Given that Gdh2 levels, but not that of PUT enzymes, decrease as the pH rises above 5, we expected that intracellular glutamate levels would increase in parallel with increasing pH given that many amino acids are continuously converted to glutamate via proline catabolism (Fig. 1B), or through specific transamination reactions [19]. We anticipated that glutamate could be converted to other amino acids to prevent the rapid and continuous generation of intracellular ammonia. For example, glutamate can be converted to glutamine by the glutamine synthetase (Gln1) enzyme that requires ammonium (NH_4_ ^+^) as a substrate [19]. We further hypothesized that the surplus of glutamate, beyond the amount that alternative metabolic pathways can assimilate it, could be excreted to the extracellular environment. Either of these processes could individually or together decrease the intracellular glutamate pool, thus relieving the toxic effects of generating ammonia. Amino acid extrusion has previously been reported to occur in *S. cerevisiae* [23], however, this has not been adequately addressed in *C. albicans.* Amino acid excretion has only been observed in studies analyzing spent growth medium from *C. albicans* biofilm cultures [24].

We measured the level of extracellular glutamate 3 h and 5 h after growth in a glutamate-devoid alkalization medium (YNB+PALAAG) with amino acids that can be metabolized to glutamate [19]; this medium contains 0.5 g/L of <u>P</u>roline, <u>A</u>rginine, <u>L</u>eucine, <u>A</u>lanine, and <u>A</u>spartate, and 1% <u>G</u>lycerol as primary carbon source. Extensive washing (4x) of the pre-cultured YPD grown cells was crucial to remove contaminating amino acids, including glutamate, prior to shifting to the YNB+PALAAG medium. We chose 3 h as the earliest time point as this correlates with the dramatic reduction in Gdh2 levels in cells growing in the unbuffered YNB+CAA medium (Fig. 1). As expected, we detected glutamate in the spent medium from the wildtype culture (Fig. 3). We also detected glutamate in the spent medium from the *gdh2-/-* mutant after 3 h, although this was significantly lower than the wildtype (*p*<0.0001). Interestingly, 2 h later (i.e., 5 h post-YPD shift), we observed a dramatic removal (∼20-fold) of extracellular glutamate in the wildtype culture, suggesting that glutamate is transported back into wildtype cells. On the contrary, extracellular glutamate increased in the *gdh2-/-* strain, indicating continued excretion. Together, glutamate extrusion appears to provide a novel mechanism by which the fungus can modulate its intracellular glutamate pool to limit ammonia production.

**Fig. 3.**
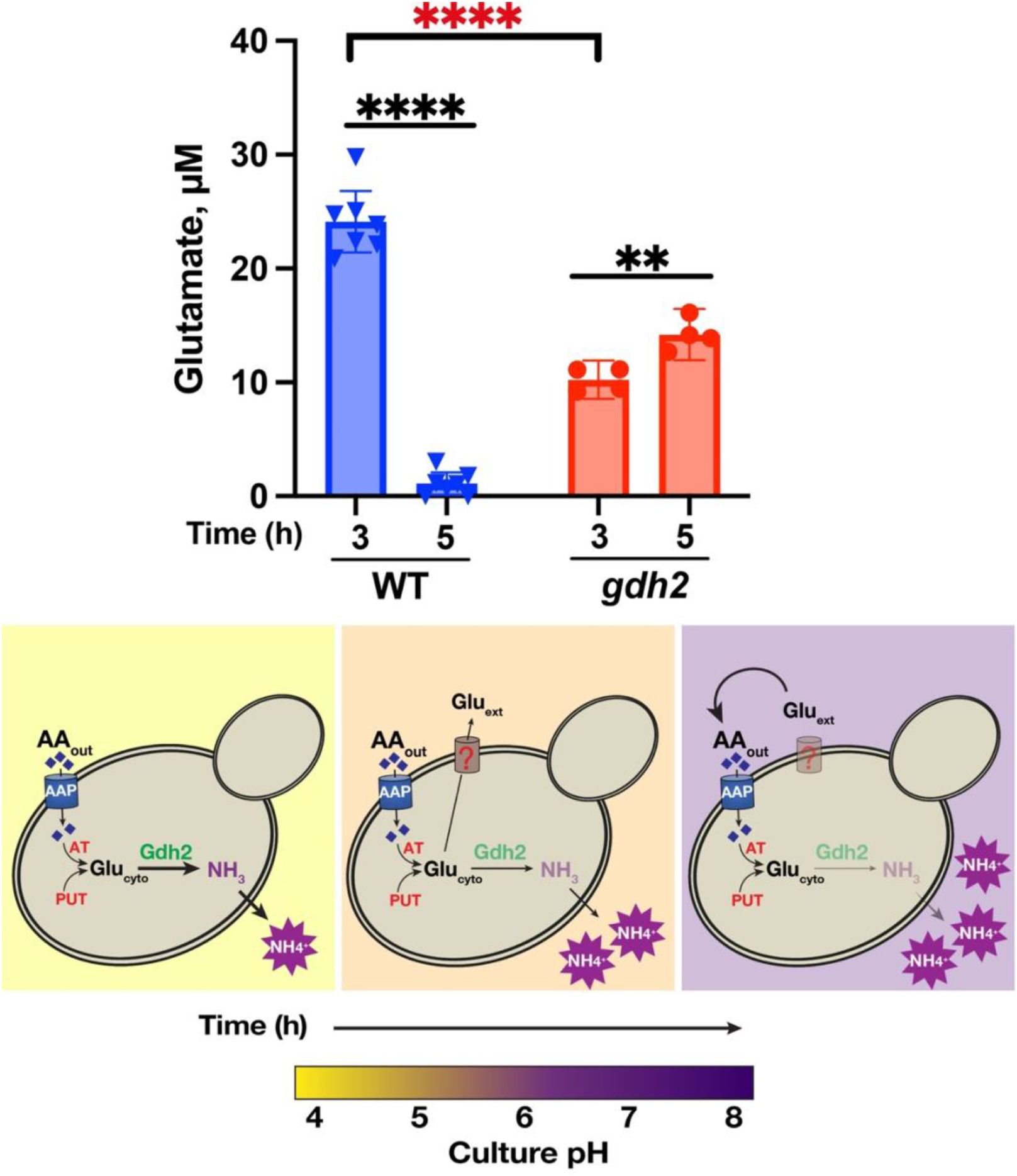
Glutamate extrusion in *C. albicans* as a mechanism to regulate intracellular glutamate pool. (Upper panel) Glutamate levels in YNB+PALAAG medium grown with WT (SC5314) and *gdh2* (CFG279) cells taken at the indicated timepoints. Data shown were obtained from seven (WT) and four (*gdh2*) biological replicates (mean with 95% CI; \*\*\*\**p* <0.0001 and \*\**p* <0.01 by student *t*-test; red asterisks denote comparison between WT and *gdh2* at 3 h timepoint). (Lower panel) Schematic diagram of glutamate extrusion in *C. albicans* as a function of time. Cytosolic glutamate (Glu_cyto_) generated from aminotransferases (AT) reactions or proline catabolism (PUT) are converted by Gdh2 to ammonia (NH_3_), which neutralizes the extracellular space. As the pH increases, Gdh2 level decreases which provide a surplus of Glu_cyto_ that are then extruded to the medium and then reassimilated.

### Stp2 and Put3 are key regulators of Gdh2, Put1, and Put2 expression

The inducible nature of Gdh2 expression during growth in a CAA-based medium motivated us to study the regulatory mechanisms driving its expression. Interestingly, Gdh2 induction is independent of its substrate (glutamate) but is dependent on proline and arginine through the proline utilization transcription factor, Put3; however, it appears that arginine provides another layer of regulatory control as the *put3* mutation failed to fully suppress arginine-dependent Gdh2 induction [17]. We rationalized that the two transcription factors (TFs) Stp2 and Put3 are key regulators of *GDH2* expression since strain lacking *STP2* (*stp2*Δ/Δ) showed a severe alkalization defect in amino acids [8] and that *PUT3* is required for the proline-dependent Gdh2 expression [17]. PathoYeastract [25] also identified Stp2 and Put3 as positive regulators of *GDH2* based on transcriptional profiling studies [11, 26]. To test this, we inactivated *STP2* or/and *PUT3* in strain CFG441 (Fig. S1A) and tested the mutant strains’ capacity to alkalinize YNB+CASG medium (Fig. 4A, upper panel). It is noteworthy to highlight the versatility of CRISPR/Cas9 by enabling the simultaneous inactivation of all three *STP2* alleles (Fig. S1A) [14]; we designate *stp2-/-/-* as *stp2*. As in previous reports, the *stp2* strain showed a clear defect in alkalization even at high density (OD ≍ 5) (Fig. 4A, upper panel). However, albeit exhibiting a clear defect, the *stp2* strain eventually alkalinized the media upon prolonged incubation (24 h). On the contrary, the *put3* strain showed a modest delay in alkalization (Fig. 4A, upper panel) but was significant as indicated by the pH of the culture measured after 24 h (Fig. 4A, lower panel). Strikingly, the *stp2 put3* double mutant showed a severe alkalization defect that clearly surpassed that of *stp2* mutant and that almost phenocopied the abolished alkalization phenotype of *gdh2* mutant (Fig. 4A, upper panel). We noted, however, that relative to YNB+CASG, the *stp2* or *stp2 put3* mutants showed an even more prominent defect in YNB+CAA after 24 h (Fig. S1B), which is likely due to the tight dependency of amino acid uptake that is dependent on Stp2. We also inactivated *PUT3* in the previously reported *stp2*Δ/Δ strain [8] and obtained identical results.

**Fig. 4.**
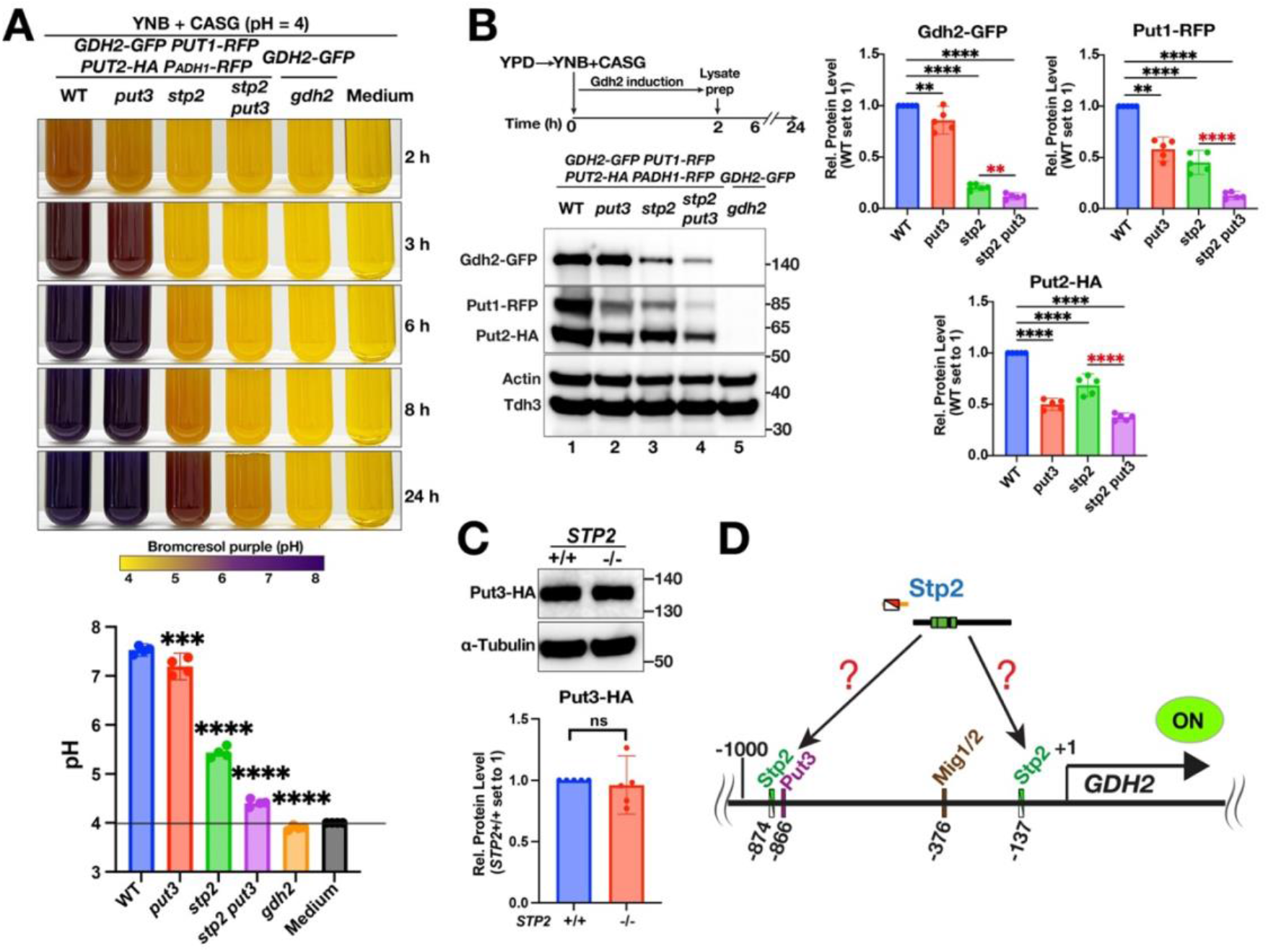
Gdh2 expression is co-regulated by Stp2 and Put3. A) Cells of the following genotypes were grown in YNB+CASG at OD ≍ 5 and then photographed at the indicated timepoints (Top) and corresponding pH after 24 h (Bottom). Strains used: WT (CFG441), *put3* (CFG443), *stp2* (CFG665), *stp2 put3* (CFG671), and *gdh2* (CFG412). **B**) Immunoblot analysis of Gdh2-GFP, Put1-RFP, and Put2-HA in the same strains as in (**A**) grown for 2 h in YNB+CASG at starting OD ≍ 2. Data are presented as Relative (Rel.) protein level (mean with 95% CI, n = 5; \*\*\*\**p* <0.0001, \*\**p* <0.01 by one way ANOVA with Dunnett’s post hoc test); Signals were first normalized to loading control (Tdh3) and then transformed the values relative to WT set to 1. Red asterisks indicate localized comparison by student *t*-test. **C**) Put3 expression is independent of Stp2. Strains CFG233 (*PUT3-HA*) and CFG676 (*PUT3-HA stp2-/-*) were grown to log phase in SGL medium and then processed for immunoblotting to detect the levels of Put3-HA. Representative immunoblot (left) and quantification (right) derived from 5 biological replicates (mean with 95% CI). **D)** Proposed model of *GDH2* regulation by Stp2. Stp2 directly regulates *GDH2* by binding directly to the upstream activating sequence (UAS) in its promoter or indirectly through regulation of an intermediate TF that directly activates *GDH2*. The presence of Stp2, Put3 and Mig1/2 binding motifs in the promoter of *GDH2* were predicted using PathoYeastract [25] and/or YeTFaSCo [27] databases.

These findings prompted us to examine the expression of Gdh2 in these strains. However, since the extracellular (alkaline) pH has a negative effect on Gdh2 levels, as shown in Fig. 1B, we limited the analysis to cultures grown for 2 h post-YPD shift from an initial OD ≍ 2. Since these strains (except for the *gdh2* control) were created in the CFG441 background, we simultaneously analyzed the levels of Put1-RFP and Put2-HA in these strains. The results clearly show that Stp2 and Put3 co-regulate Gdh2 expression, with Stp2 having a more dominant role in its regulation compared to Put3 (Fig. 4B). Interestingly, Stp2 also regulate Put1 and Put2 as indicated by their reduced levels in a *stp2* strain with Put1 being slightly more under the control of Stp2 than Put3 whereas the opposite applies for Put2. This effect does not occur indirectly through Put3, as Put3 levels were not altered in a *stp2* mutant (Fig. 4C). In all cases, the difference in the expression between *stp2* and *stp2 put3* are always significant (Fig. 4B, red asterisks) suggesting that Stp2 and Put3 operate independently and in parallel to regulate the expression of *GDH2*, *PUT1*, and *PUT2*.

Sequence analysis of the –1000 bp region preceding the *GDH2* ORF using the PathoYeastract [25] and/or YeTFaSCo [27] databases indicate the presence of a putative Put3-binding motif [CGG(Nx)CCG] [28] positioned at –866 from the ORF. Analysis of the same sequence stretch for Stp2 binding motifs [CGGCTC [29] and/or GYGCCGYR [30]; 1 bp substitution allowed] found positions –137 and/or –874 as putative binding sites in *GDH2.* Interestingly, Stp2 is not in the list of transcription factors associated with either *PUT1* or *PUT2* regulation, although we found positions –153/-668 (*PUT1*) and –197 (*PUT2*) resembling Stp2 binding motifs [29, 30]. Further analyses of these promoters indicated the presence of bindings motifs for Mig1 and Mig2 [*GDH2* (–376), *PUT1* (–165, –651, –722, –937), and *PUT2* (–94, –163)], which are partially redundant orthologs mediating glucose repression in *C. albicans* [31]. This is consistent with our previous findings that these enzymes are sensitive to glucose repression [5, 6]. In summary, the results strongly suggest that the prominent alkalization defect observed in the *stp2* mutant is primarily due to its inability to induce *GDH2* expression fully and also of the two mitochondrial-localized enzymes, Put1 and Put2, in a manner that is independent of the control of the previously characterized transcription factor Put3 [5, 26].

### The role of Gdh2 is conserved in other pathogenic *Candida* species

The capacity to neutralize the environment in the presence of amino acids was reported previously in other members of the pathogenic *Candida* species complex [2], including the multi-drug resistant *C. auris* [32] and *C. glabrata* [33]. To test whether Gdh2-dependent alkalization is conserved in these species, we created *GDH2* deletions in *C. auris* and *C. glabrata* by homology-directed recombination using an HA-tagging cassette (pFA6a-3HA-SAT-flipper) derived from the original SAT1-flipper cassette [34]. We created reconstituted strains following the procedure described previously in *C. albicans* [6] (Fig. 5A). We compared the alkalization capacity of each strain grown in dense cultures (OD ≍ 5). Interestingly and consistent with a previous report [2], *C. glabrata* does not readily alkalinize the extracellular medium (YNB+CASG) even after 72 h of prolonged growth (Fig. 5B). However, when we grew the wildtype or reconstituted strain (*gdh2*Δ*::GDH2*) in YNB+CAA where amino acids are used as sole carbon/nitrogen/energy source, alkalization becomes evident after ∼5-6 h of growth, which becomes more pronounced after 8 h (Fig. 5B). As expected, the Cg*gdh2* strain is completely unable to alkalinize the medium even after prolonged incubation (32 h). Unlike *C. glabrata*, the wildtype *C. auris* strain robustly neutralized the YNB+CASG medium, which completes after ∼5-6 h. As expected, the cultures grown with Cau*gdh2* remained acidic even beyond 24 h (Fig. 5C). The results suggest that consistent with the role of Gdh2 in *C. albicans* (CaGdh2), the Gdh2 orthologues in *C. auris* (CauGdh2) and *C. glabrata* (CgGdh2) are also essential for amino acid-dependent alkalization in these species. Analysis of the N-termini of CauGdh2 (B9j08_004192p) or CgGdh2 (CAGL0G05698g) by MitoFates [35] revealed no mitochondrial presequence (probability ∼0.00) suggesting that these enzymes, as in *C. albicans*, are also localized extra-mitochondrially in the respective species.

**Fig. 5.**
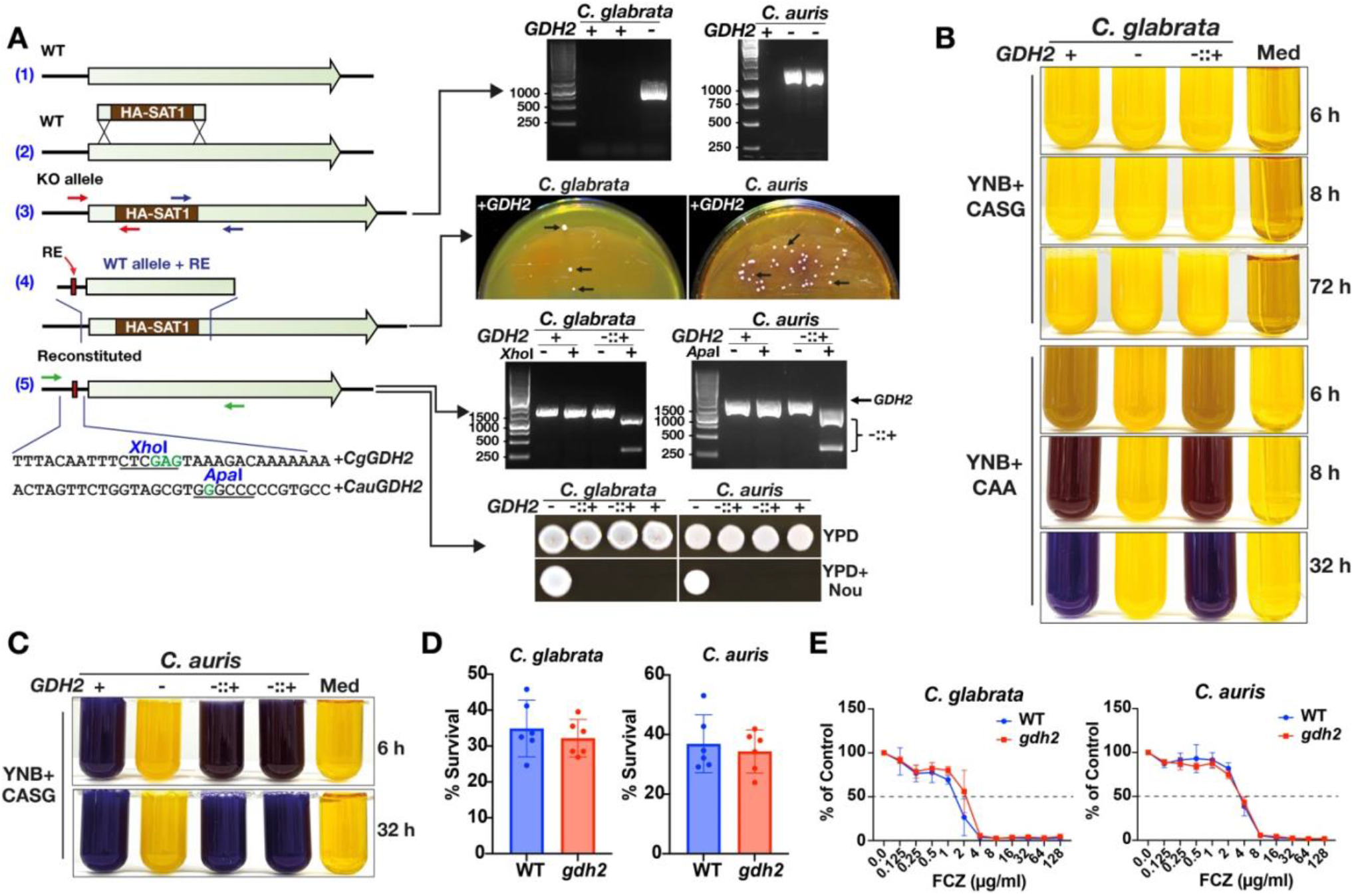
Gdh2 function is conserved in other *Candida* species. A) Genetic construction of *gdh2* and reconstituted *C. glabrata* and *C. auris* strains. A region of the wildtype (WT) gene (1) was deleted by homology-directed recombination using a cassette amplified from a plasmid derivative of the SAT1-flipper cassette (2) and correct integration was verified by PCR using one of the primer pairs indicated in the scheme (3). The corresponding gels on the right used the primer pairs in blue. For reconstitution, the deleted region was replaced by WT allele bearing restriction sites (RE) (4) that were inserted in the primers (5, green font). Transformation was made on YNB+CAA plate and colonies pointed with arrows (4) were analyzed by PCR and restriction digest using either *Xho*I (*C. glabrata*) or *Apa*I (*C. auris*) (5). The reconstituted strains lost the nourseothricin resistance marker as indicated by the lack of growth on YPD with 200 μg/ml nourseothricin (200N). **B, C)** Wildtype (+), *gdh2* (–), and reconstituted (–::+) strains of *C. glabrata* (**B**) and *C. auris* (**C**) were grown in the indicated alkalization medium at OD ≍ 5 and photographed at the indicated timepoints. **D)** *GDH2* is dispensable for survival of *C. glabrata* and *C. auris* in a human blood infection model. Cells (∼10^5^ CFU) were added into a blood aliquot and survival was assessed 1h after by plating. Data shown were obtained from six independent donors (mean with 95% CI; not significant by student *t*-test). **E)** *GDH2* is not required for fluconazole (FCZ) susceptibility of *C. glabrata* and *C. auris*. Growth (as OD) of the indicated strains in increasing concentrations of fluconazole was measured at 24 h. Growth inhibition of 50% (MIC) relative to untreated control is shown as a dotted horizontal line. Each data point represents mean ±SD (n=3-4). Strains used: *C. glabrata* – wildtype (+, ATCC 2001), *gdh2* (–, CFG670), reconstituted (–::+, CFG679); *C. auris* – wildtype (+, CFG552), *gdh2* (–, CFG586), reconstituted (–::+, CFG699)

Despite its central role in amino acid metabolism, Gdh2 is dispensable for the virulence of *C. albicans* [6]. To assess the role of Gdh2 in virulence of *C. auris* and *C. glabrata*, we assayed the survival of the wildtype and *gdh2* strains using a human whole blood infection model [36]. Consistent with our findings in *C. albicans*, Gdh2 is dispensable for virulence of these two *Candida* species as there was no significant difference in the survival of *gdh2* mutant in either species relative to their respective wildtypes (Fig. 6D). In addition, despite amino acid metabolism being implicated in antifungal tolerance [37], the loss of *GDH2* did not affect the fluconazole susceptibility of either *C. auris* or *C. glabrata* (Fig. 6E). In summary, although Gdh2 is the key enzyme that endows *Candida* spp. the capacity to neutralize the environment when grown in amino acids, it remains dispensable for properties associated with virulence characteristics of these species.

**Fig. 6.**
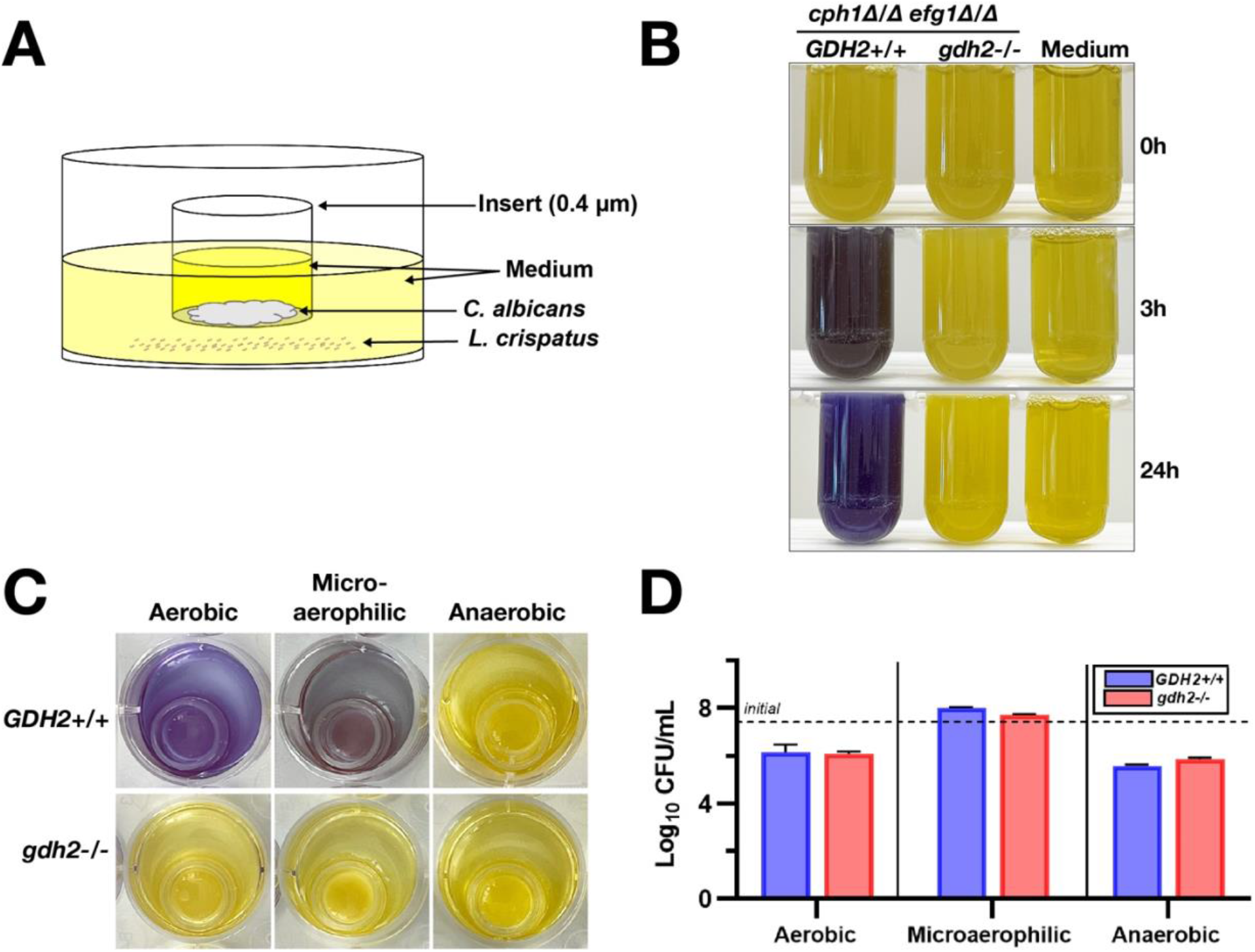
Environmental alkalization does not antagonize growth of competing microorganism. A) Schematic diagram showing the in vitro set-up to investigate the effect of pH modulation in growth of *L. crispatus*. **B)** *cph1*Δ/Δ *efg1*Δ/Δ *GDH2*+/+ (CASJ041) and *cph1*Δ/Δ *efg1*Δ/Δ *gdh2*-/-(CFG352) were grown in CNY alkalization medium at OD ≍ 5 and photographed at the indicated timepoints. **C)** Alkalization is dependent on oxygen tension. Photographs of the wells after 48 h of growth under aerobic, microaerophilic, and anaerobic growth conditions. **D)** Fungal-driven pH modulation does not antagonize *L. crispatus*. Viable cell count of *L. crispatus* recovered in the medium indicated in (**C**); Results (n=4) between *GDH2+/+* and *gdh2-/-* in each oxygen tension condition were not significant by student *t*-test.

### Environmental alkalization does not antagonize growth of *Lactobacillus*

The intriguing finding that inactivation *GDH2* does not affect *C. albicans* virulence [5] motivated us to question whether environmental pH modulation by *C. albicans* could affect the composition of the host commensal microbiome. In line with its opportunistic character, we posited that *C. albicans*-dependent alkalization of the growth environment could create imbalances in the host microflora antagonizing the growth of other competing, and potentially antagonistic microorganisms. For example, *Lactobacillus* species normally inhabit the acidic vaginal microenvironment in most healthy females, and these bacteria are thought to provide protection against *C. albicans* [38]. Acute vulvovaginal candidiasis (VVC) occurs in most women (≥75%) at least once in their lifetime and VVC represents a prime example of dysbiosis within a complex host niche [39]. Among the *Lactobacillus* species, *L. crispatus* is one of the predominant species in the vaginal tract [40] and is thought to exert the most potent antimicrobial activity against *C. albicans* [38], making it an ideal representative of this genus. To experimentally address the possibility that *C. albicans*-dependent alkalization inhibits the proliferation of *L. crispatus* we used a transwell (0.4 μm) culture system schematically depicted in Fig. 6A. We used a modified alkalization medium (CNY; see Methods) containing ∼5.6 mM (0.1%) glucose, which is comparable to the level observed in vaginal secretions [41]. Under these conditions, CNY supports the growth of *L. crispatus* and *C. albicans*. However, under aerobic conditions, *L. crispatus* fails to grow without *C. albicans* growing in the transwells; *C. albicans* apparently depletes oxygen in the media, a requisite for *L. crispatus* growth (Fig. S3). We used non-filamenting *C. albicans* strains carrying null alleles of *CPH1* and *EFG1* (*cph1*Δ/Δ *efg1*Δ/Δ) to exclude potential inhibitory factors associated with filamentation. The *C. albicans* strains were seeded into the transwell at a relatively high starting cell density (∼1.6 x 10^7^ CFU/transwell) to minimize growth-dependent effects. As expected, in contrast to the *cph1*Δ/Δ *efg1*Δ/Δ strain, the strain lacking *GDH2* (*cph1*Δ/Δ *efg1*Δ/Δ *gdh2*) was unable to alkalinize the media (Fig. 6B)[6]. The tight dependency of alkalization on respiration was clear (Fig. 6C); under aerobic and microaerophilic conditions the medium became alkaline in a Gdh2-dependent manner, whereas consistent with the requirement of mitochondrial function, alkalization did not occur under anaerobic conditions. Contrary to our expectations and despite obvious difference in extracellular pH, there were no significant differences in the number of *L. crispatus* cells recovered in the wells with transwells seeded with either *GDH2* or *gdh2 C. albicans* strains (Fig. 6D). Also unexpectedly, *L. crispatus* failed to counteract the alkalization exerted by *GDH2* strain grown under aerobic and microaerophilic conditions, which is surprising given that lactobacilli are thought to play a prominent role in maintaining the acidic vaginal pH [42] and that *L. crispatus* inhibits growth of *C. albicans* [38]. Taken together, these results clearly suggest that active pH modulation exerted by *C. albicans* may not effectively antagonize the growth of competing microorganism.

## Discussion

The interest in studying environmental alkalization as a consequence of amino acid metabolism in *C. albicans* is due its purported link in facilitating virulent growth (reviewed in: [1, 43, 44]). A precise understanding of the mechanisms underlying alkalization has remained elusive due to the lack of information regarding how ammonia is generated. *C. albicans* lacks the enzyme urease, which in many pathogenic microorganisms catalyzes the breakdown of urea to ammonia and carbon dioxide [45]. Consequently, urea amidolyase (Dur1,2), which catalyzes the analogous reaction in *C. albicans*, but in two steps, was initially considered to be a primary source of ammonia. However, the inactivation of *DUR1,2* was found to have a modest effect on alkalization [2, 5]. We recently discovered that the cytoplasmic glutamate dehydrogenase (CaGdh2) is the key enzyme catalyzing the release of ammonia and functions as the major driver of environmental alkalization [6]. With this knowledge, we have been able to assess the contribution of other peripheral factors that impinge on this process and have defined several control points. Based on our findings we have developed an integrative model that accounts for how the catabolism of amino acids leads to environmental alkalization (Fig. 7). Despite its role in alkalization, *GDH2* is dispensable for virulence of *C. albicans* in *Drosophila* and murine models of candidemia [6] and in human whole blood (unpublished data). Our results challenge several commonly accepted dogmas regarding the role of alkalization in fungal virulence.

**Fig. 7.**
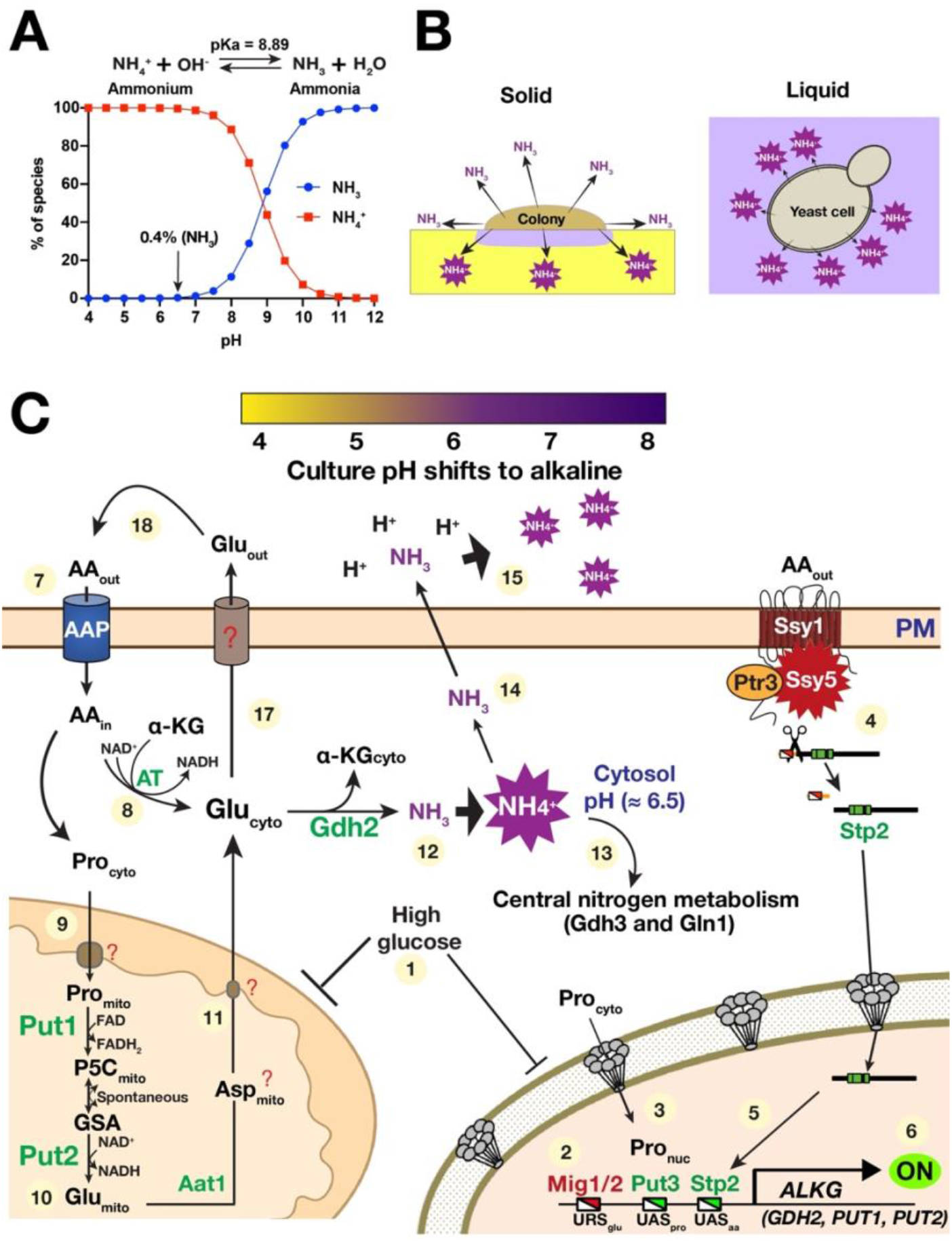
Integrative model of amino acid-dependent alkalization of the extracellular growth environment by *C. albicans*. A) A plot showing the relative concentrations of ammonia (NH_3_) and ammonium (NH_4_ ^+^) in aqueous solution based on pH set at 37 °C. Values were calculated using pKa = 8.89 [58]. The ratio of NH ^+^ to NH_3_ in this equilibrium is highly pH-dependent. **B)** Schematic diagram showing the difference in the ammonia release in liquid and solid media. In comparison to solid media, volatile ammonia is more efficiently captured by the acidic liquid culture to form ammonium resulting in a more rapid alkalization of the media. **C**) Regulatory control of alkalization gene expression and glutamate-dependent metabolism determines the alkalization potential of *C. albicans*. When cells are shifted from high glucose medium (e.g., YPD) to alkalization medium (e.g., YNB+CAA), *C. albicans* cells are relieved of glucose repression (1) transcriptionally mediated by Mig1/2 repressors binding upstream repressing sequences (URS_glu_)(2). Cytosolic proline (Pro_cyto_) enters the nucleus (Pro_nuc_) where it binds and activates the transcription factor Put3 at upstream activating sequence (UAS_pro_) (3). In parallel, the transcription factor Stp2 is proteolytically activated by the SPS sensor (Ssy1-Ptr3-Ssy5) in response to extracellular amino acids (4). The processed form of Stp2 efficiently targets the nucleus and binds UAS_aa_ (5). The coordinated activities of these factors determine the transcriptional output of the key alkalization genes (6). Uptake of extracellular amino acids is facilitated by amino acid permeases (AAP) (7). Many amino acids are converted to glutamate in the cytosol (Glu_cyto_) by transamination reactions (8) that require α-ketoglutarate (α-KG) as a substrate. Proline, enters the mitochondria via an unidentified (?) transporter (9) where it is converted by the proline catabolic pathway (PUT1 and PUT2) to glutamate (Glu_mito_) (10). It is postulated that Glu_mito_ is directly or indirectly (?) exported out of the mitochondria (11) and contributes to the Glu_cyto_ pool. This multi-step path involves conversion of Glu_mito_ to aspartate in the mitochondria (Asp_mito_), predicted to be catalyzed by the mitochondrial aspartate aminotransferase (Aat1). Asp is exported to the cytosol where it gets converted to Glu_cyto_ by cytosolic aspartate aminotransferase. Glu_cyto_ is the substrate of Gdh2 forming α-KG and ammonia (NH_3_) (12). Due to the cytosolic pH, maintained at around 6.5, the bulk of ammonia rapidly converts to ammonium (NH ^+^). Ammonium is a substrate of central nitrogen metabolism (Gdh3 and Gln1) (13). NH, although a minor species, can efficiently diffuse across the cell membrane (14) where it forms NH_4_ ^+^ and thereby effectively contributes to neutralizing the extracellular space (15). As the extracellular pH increases, Gdh2 level decrease, and a fraction of Glu_cyto_ is extruded transiently (Glu_ext_) by a still uncharacterized (?) exporter (17). Glu_ext_ is reassimilated after transport into cells via AAP (18).

Based on the pK_a_ of ammonium/ammonia (NH ^+^/NH) relationship in aqueous medium (Fig. 7A), the ammonia generated by Gdh2 in the cytosol (pH ∼ 6.5) is expected to rapidly form ammonium. Ammonium can be reassimilated by NADPH-dependent glutamate dehydrogenase (Gdh3; α-ketoglutarate to glutamate) and/or glutamine synthetase (Gln1; glutamate to glutamine)[19]. However, it is unlikely that these anabolic reactions can accommodate the amount of excess ammonium generated when cells are metabolizing amino acids as energy sources. Rather, our data are consistent with the notion that ammonia, although present in small amounts (∼0.4%), passively diffuses across the cell membrane into the extracellular environment where it contributes to neutralizing the acidic pH as it forms ammonium. This occurs without needing a dedicated exporter [46–48]. This fully accounts for the observed density-dependent alkalization; higher cell densities result in more ammonia released and faster alkalization. Also, this explains why liquid cultures rather than colonies growing on solid substrates, exhibit more rapid alkalization. The volatile ammonia (NH_3_) formed by colonies diffuses in multiple directions limiting the amount that is directly captured by the solid growth medium (Fig. 7B). Alkalization in liquid cultures occurs faster, and when coupled to a high cell density, the pH changes rapidly. We tested the role of Ato5, a putative ammonia transport protein, by creating an *ato5-/-* strain. Contrary to previous reports [2, 16], we were unable to confirm a major effect on alkalization (Fig. S2A, S2B); thus, we excluded Ato5 from our model. When assayed in liquid culture, we can reliably link the alkalization deficiency of a given strain to an intrinsic critical factor driving alkalization in a manner that is less dependent on growth, e.g., the *nuo1*Δ/Δ strain (Fig. 2E). In fact, the ability of a null mutant strain to abrogate alkalization of a dense liquid culture (OD ≍ 5) is an “acid test” for being an essential component of alkalization. To our knowledge, Gdh2 is the only component that clearly passes this test; i.e., strains lacking Gdh2 activity fail to alkalinize the growth media.

Although Gdh2 is a cytosolic component [6], our model accounts for the tight dependence on mitochondrial function (Fig. 7C). The substrate glutamate originates mainly from metabolic events in the mitochondria (Fig. 2B, 2C). Consistently, conditions that downregulate mitochondrial function, such as growth in high glucose (2%), limiting oxygen (Fig. 2F, 5C), the lack of mitochondrial respiratory subunits (*nuo1*Δ/Δ), or inhibition by Antimycin A, all result in alkalization deficiency. Indeed, regardless of how amino acids are used by the cells, i.e., as sole carbon/nitrogen/energy source (YNB+CAA), as sole nitrogen source (YNB+CAA supplemented with 1% glycerol, 1% lactate, or <0.2% glucose as main carbon source), or as a supplementary carbon/nitrogen/energy source (YNB+CASG), alkalization readily occurs as long as conditions permit respiratory growth. High glucose represses mitochondrial function and Put1 and Put2 expression, thus lowering intracellular glutamate (Fig. 2A). The coupling of reduced intracellular glutamate with the downregulation of mitochondrial function explains why alkalization is virtually abolished in high glucose. Strains lacking *PUT1* and *PUT2* exhibit clearly delayed alkalization even in dense cultures [5, 6], but alkalization is ultimately achieved presumably due to multiple transamination reactions that generate glutamate [19]. We note that high glucose-mediated repression of the mitochondrial function can result in the generation of fermentation by-products and CO_2_, which contribute to maintaining acidic pH [5, 49].

We traced the prominent alkalization defect of strains carrying *stp2* to the inability to fully derepress *GDH2*, *PUT1*, and *PUT2* (Fig. 4). This is consistent with, but adds to our recent findings [17] that the proline-dependent expression of Put1, Put2, and Gdh2 is partially independent of Put3. The additive effect of combining *stp2* and *put3* mutations indicate that Stp2 and Put3 function in parallel to regulate the enzymes contributing to alkalization in *C. albicans*. This relationship explains some of the interesting *stp2* phenotypes that are relevant to *C. albicans* pathogenesis, such as the defective filamentation in the phagosomes of macrophages [8] and virulence in murine systemic infection model, which we have shown to be dependent on proline catabolism [5, 17]. This also helps explain other findings related to the increase in proline uptake in *stp2* mutant [24], which is likely a compensatory mechanism to generate more energy when intracellular amino acids become limiting. The tight dependency of Gdh2 expression on Stp2 presented an apparent conundrum given that Gdh2 expression is extremely low in YPD [6] even though Stp2 is constitutively expressed and activated [10, 14]. However, this can be understood since *GDH2* expression is apparently subject to glucose repression by Mig1 and Mig2; these factors have been shown to negatively regulate *GDH2*, *PUT1,* and *PUT2* [31]. Shifting cells from YPD to the alkalization medium with low glucose relieves the repression, presumably enabling Stp2 and Put3 to activate expression by binding to putative UAS_aa_ and UAS_Put3_ in the promoters of target genes, respectively. Further work is required to experimentally define the precise nature of the promoter regions in these genes. For example, since Gdh2 is weakly expressed in YNB+CAA with 2% glucose (Fig. 2B), glucose repression cannot fully explain the lack of Gdh2 expression in YPD grown cells [6].

The observation that Gdh2 levels decrease as the extracellular pH increases and cells extrude glutamate suggests that cells do this to limit ammonia production and thereby actively respond by reducing the risk of creating an alkaline growth environment. A similar strategy has been demonstrated in *S. cerevisiae*, which limits the production of ammonia by excreting amino acids, including glutamate via transporters that belong to the multidrug resistance transporter family postulated to function as H^+^ antiporters (e.g., Aqr1) [23]. We note that the *C. albicans* genome encodes a putative *AQR1* homolog (*QDR2*/C3_05570W), however we have not investigated if it operates similarly to *S. cerevisiae*. We found that the extrusion of glutamate by *C. albicans* is transient; the excreted glutamate is eventually reassimilated. It is unclear whether other amino acids are also excreted in addition to glutamate, but it would not be surprising if they are. The fact that *C. albicans* extrudes glutamate, which reduces the alkalization potential, suggests that as other fungi, they prefer acidic growth conditions that provide a competitive advantage over many bacteria, is more favorable to facilitating nutrient uptake powered by the proton gradient, and minimizes the potential loss of nitrogen. Finally, maintaining an acidic growth environment likely contributes to keeping intracellular ammonium levels below toxic levels. Ammonia, not ammonium, readily diffuses across the plasma membrane [46–48] in a direction towards a lower pH, effectively pulling the ammonium/ammonia equilibrium to drive excess ammonium out of cells.

As in *C. albicans*, Gdh2 is essential for the growth of *C. glabrata* and *C*. *auris* in media with amino acids as the sole carbon/nitrogen/energy source (YNB+CAA). We exploited this phenotype during strain constructions (Fig. 5A). Consistent with our finding that Gdh2 is dispensable for *C. albicans* virulence [6], *GDH2* appears also to be dispensable for virulence of both *C. glabrata* and *C. auris*, as assessed by survival in a whole human blood. These results indicate that the other alternative carbon sources suffice for growth of *gdh2* strains in these infection models. Our future efforts will examine how *Candida* cells adjust their metabolism in the context of infection, which we posit involves transcriptional and/or translational arrest; hence our current work includes the use of high-throughput 5′P degradome RNA sequencing (HT-5Pseq) to investigate the degradation of 5′phosphorylated mRNA intermediates as a readout for 5′-3′ co-translational mRNA decay and ribosome stall [50].

Our observation that the fungal-dependent pH modulation also does not influence the growth of *L. crispatus*, a representative of competing microorganism (Fig. 6), supports frequent findings of vaginal yeast colonization within *Lactobacillus*-dominant vaginal microbiomes [51]. It suggests that *C. albicans* itself does not exert a direct inhibitory role towards lactobacilli to alter the microbial population of the vaginal tract. Rather, external factors aside from pH might contribute to their overgrowth, especially in conditions that allow easier access to host epithelial cells [52]. One theory is that this alkalization may contribute to dysbiotic consequences that reduce lactobacilli. High vaginal pH may create microbiome shifts favorable to disease-associated vaginal microbes and open up niches that can serve as opportunities for *Candida* overgrowth [53]. In the transwell assay, it was striking to see that under aerobic and microaerophilic conditions, *L. crispatus* was unable to counteract the alkalization by *C. albicans* in a culture medium containing low but physiological levels of glucose (0.1%)[41]. Although this experimental set-up does not take into account alternative carbon sources, it clearly shows that to counteract the alkalization potential of Candida, *Lactobacillus* sp. must effectively ferment other carbohydrate sources present in the vaginal milieu (e.g., glycogen) to maintain the acidic vaginal pH [38]. Interestingly, clinical observations show that vaginitis due to *Candida* overgrowth, in contrast to that caused by bacteria or *Trichomonas*, is not associated with an increase in vaginal pH [54]. Together with our data, this shows that the strict dependency of fungal-driven alkalization on respiration and oxygen tension (Fig. 2F, 6C) is consistent with the notion that the vaginal microenvironment is oxygen-limiting. Even if there is an increase in oxygen tension in the vagina due to a variety of factors, such as increased sexual activity and menstruation [55], the outcome will still result in the vaginal pH remaining acidic, suggesting that other unexplored factors might still contribute to the repression of fungal-dependent alkalization. Our results indicate that the inhibitory effects of *L. crispatus* and other *Lactobacillus* species towards *C. albicans* do not involve mitochondrial repression in a manner analogous to the effect of phenazine produced by *Pseudomonas aeroginosa* [49]. Consistently, *Lactobacillus*-mediated inhibition of *C. albicans* hyphal formation was proposed to occur via secretion of the small molecule 1-acetyl-β-carboline (1-ABC) that inhibits Yak1, a member of the dual-specificity tyrosine phosphorylation-regulated kinase (DYRK) that is predicted to localize in the nucleus and cytoplasm as that reported in *S. cerevisiae* [56]. Although our data suggest that active pH modulation is not directly required for virulence of *C. albicans* and other *Candida* spp., the results warrant the reassessment of virulence processes reportedly linked to or associated with pH modulation by fungal pathogens.

## METHODOLOGY

### Organisms and culture

All key materials (organisms, chemicals, oligonucleotides, and software) used in this work are summarized in Table S1 (Key resource table). Yeast strains were routinely cultivated on YPD agar (1% yeast extract, 2% peptone, 2% glucose, and 2% Bacto agar) following recovery from glycerol stock stored in a –80 °C freezer. Where needed, YPD is supplemented with nourseothricin (Nou) at the required concentrations (200-, 100-, and 25-μg/ml) from filtered stock (200 mg/ml). Overnight YPD broth cultures were prepared by picking single colonies from a YPD plate and grew in a shaking incubator (Infors HT Multitron Incubator Shaker) set at 30 °C and 150-180 rpm. Alkalization was routinely assessed using the standard YNB+CA medium (0.17% yeast nitrogen base without amino acids and ammonium sulfate (YNB), 1% of casamino acids (CA) containing 0.01% Bromocresol Purple (BCP) as pH indicator. This medium and other alkalization media were adjusted to pH = 4.0 using 1 M HCl before sterile-filtered (0.45 μm). The base medium was used without or with 2% glucose depending on the experiment. Where indicated, YNB+CAA is supplemented with 38 mM ammonium sulfate (5 g/L ≍ 38 mM) and 1% glycerol (YNB+CASG) to support the growth of some mutants. For glutamate extrusion analysis, YNB was supplemented with 0.5 g/L each of <u>P</u>roline, <u>A</u>rginine, <u>L</u>eucine, <u>A</u>lanine, and <u>A</u>spartate, and added with 1% <u>G</u>lycerol as the main carbon source, which is referred elsewhere in the text as YNB+PALAAG medium and used without BCP. For the alkalization experiment with *Lactobacillus crispatus*, the CNY medium was composed of YNB (0.085%), CAA (0.5%), proteose peptone no. 3 (0.375%), yeast extract (0.095%), NaCl (0.125%), glucose (0.1%), BCP (0.01%); this medium is a 50:50 mix of standard YNB+CAA medium and NYCIII medium (without glucose) and then supplemented with glucose and BCP at the indicated concentrations. Other specific media modifications are indicated elsewhere in the text. Stock solutions of different carbon and nitrogen sources used are as follows: glucose (40%), glycerol (20%), and ammonium sulfate (1 M).

### CRISPR/Cas9 gene editing

For inactivation of *STP2*, the pV1524 vector bearing the *STP2* 20-bp sgRNA (pFS136, derived from primers p6/p7), repair templates (RT) with in-frame stop codon and *Xho*I restriction site (p8/p9), and verification primers (p10/p11) can be found in Table S1. For *ATO5* inactivation, two sgRNAs were designed corresponding to pFS141 (p12/p13) and pFS143 (p14/p15). Since these sgRNA regions are in close proximity, a common set of RT (p16/p17) and verification primers (p18/p19) were used for both. The *PUT3* vector (pFS084), RT (p2/p3), and verification primers (p4/p5) were previously described [5]. Specific sgRNAs were cloned in either pV1093 (*PUT3*) and/or pV1524 (*STP2 or ATO5*) by blunt-end ligation. Cassettes were verified by sequencing using primer p1 and then released by digestion with *Kpn*I/*Sac*I (5 μg plasmid/digestion). Digested cassettes and RTs (generated by template-less PCR) were PCR-purified and co-transformed into the indicated strain using the hybrid lithium acetate/DTT-electroporation method, as previously described in [5]. Nourseothricin-resistant (Nou^R^) transformants were selected on YPD+200 μg/ml Nou, and further clonal purification was made on YPD+100 μg/ml Nou. Transformants were purified and verified several by colony PCR using the appropriate primers to amplify the mutated gene and then digested with *Xho*I to identify the knockouts. For pV1524-derived plasmids, cassette excision was done by growing purified colonies in liquid YPM and then plating on YPD+25 μg/ml Nou. Nourseothricin-sensitive (Nou^S^) pop-outs were verified by streaking on YPD+100 μg/ml Nou. For the construction of *stp2-/– put3-/-* double knockout strain, the Nou^s^ *stp2-/-* strains were co-transformed with the digested pFS084 and RT, and transformants were selected accordingly. In most cases, single preparations of purified digested cassettes and RT sufficed for 10-15 independent transformations and were kept at –20 °C until used.

### Genetic construction of *gdh2* and reconstituted *C. glabrata* and *C. auris* strains

Since *C. glabrata* and *C. auris* are haploids, a homology-directed recombination gene deletion strategy was used to delete *GDH2* in these strains. Briefly, a knockout cassette was amplified from an HA-tagging plasmid (pFS069 = pFA6a-3HA-SAT flipper) [5] using primer pairs p21/p22 for *C. glabrata GDH2* (CAGL0G05698g) and p28/p29 for *C. auris GDH2* (B9J08_004192). pFS069 was derived from the original *SAT1*-flipper cassette [34]. This tagging cassette has a very good recombination efficiency even if the homology flanking regions to the target genes are short (<100-bp). Amplified cassettes were purified and transformed into *C. glabrata* (ATCC 2001) and *C. auris* (CFG552) wildtype strains and Nou^R^ transformants selected on YPD+200 μg/ml Nou. Purified colonies were checked for correct integration, and verified knockouts were stored at –80 °C as glycerol stock without removing the NAT^R^ marker, as it would be used as a marker for reconstitution. The following primers were used to verify junctions for correct knockout cassette integration: *Cg*-*gdh2*Δ (p27/p24; p23/p34) and *Cau*-*gdh2*Δ (p27/p30; p33/p34). For genetic reconstitution of knockout strains, a region of the *GDH2* gene was amplified by PCR from the respective wildtype strains using primers p25/p26 for *C. glabrata GDH2* and p31/p32 for *C. auris GDH2*. The amplicons were purified and then transformed into the respective knockout strains. Electroporated cells were recovered in YNB+CA medium for 6-8 h and then plated on YNB+CA agar. Plates were incubated for 2-3 days at 30 °C until colonies appeared. These colonies were purified on YPD, and then 3-5 independent colonies per transformation were tested for correct integration by PCR (*C. glabrata*, p23/p24; *C. auris*, p33/p30) and restriction digest using either *Xho*I (*C. glabrata*) or *Apa*I (*C. auris*). The verified reconstituted strains were tested for growth on YPD agar with 200 μg/ml nourseothricin to determine the loss of the Nou^R^ marker, indicating replacement by the reconstituted allele.

### Alkalization assay

For routine assays, single colonies from YPD plates were picked and grown overnight (16-20 h) in YPD broth and then collected the following day by centrifugation at 4000 *g* for 3-5 min. Depending on the number of cells needed, overnight cultures were prepared in 3-, 25– or 50 ml volumes. Harvested cells were washed twice with ddH_2_O and then resuspended in ∼20% of the original culture volume to concentrate the cells. Optical density (OD) at 600 nm was measured, and used this value to inoculate prewarmed YNB+CAA or YNB+CASG medium (37 °C) at OD ≍ 0.1, 2, or 5. Where indicated, YNB+CAA medium was supplemented with 2% glucose. Cultures were incubated in a 37 °C shaking incubator (150-180 rpm), and depending on the experiment and specific timepoints, cultures were photographed and/or sampled for further analysis.

### Enzyme expression analysis

A one ml aliquot of the culture at 2 h timepoint prepared from a starting OD ≍ 2 was immediately mixed with 250 μl of cold 2 M NaOH with 7% β –mercaptoethanol (βME) to lyse the cells. After 15 min, an equal volume of 50% trichloroacetic acid (TCA) was added to precipitate the proteins, which was carried out overnight at 4 °C. Protein pellets were harvested by high-speed centrifugation (17,000 *g*, 10 min, 4°C) and then the residual liquid in the tube completely removed by a quick high-speed spin. Protein precipitates were solubilized completely in a 50 μl 2X SDS sample buffer containing 5% βME and 167 mM of Tris Base (pH ≍ 11) and then boiled at 95°C for 5 min. Proteins were resolved in 4-12% Bis-Tris pre-cast gel using either MES or MOPS buffer and then subjected to a standard immunoblotting procedure in a nylon membrane. After transfer, membranes were blocked using 10% skimmed milk in TBST (TBS + 0.1% Tween) for 1 h at room temperature. Target proteins were detected individually or simultaneously using an optimized antibody cocktail prepared in 5% skimmed milk in TBST for both primary (α-GFP (1:3000), α-mCherry (1:6000), α-Tdh3 (1:5000), α-actin (1:5000,) α-tubulin-HRP (1:10000)), and secondary (goat α-mouse poly-HRP (1:15000), goat α-rabbit poly-HRP (1:15000), α-HA-HRP (1:15000)). When used in a cocktail, α-tubulin-HRP (1:10000) is added to the secondary antibody as it is already conjugated to HRP. A chemiluminescent substrate (SuperSignal Dura West Extended Duration Substrate) was added to detect immunoreactive bands in blots using the Azure detection system.

### Cycloheximide (CHX) treatment

Cells collected from YPD cultures were inoculated into 20 ml of YNB+CA at OD ≍ 2 and then grown continuously with shaking at 37 °C. After 2 h, 3 ml culture aliquots were added to separate tubes containing concentrated buffers (500 mM, pH = 4-8) diluted by cultures to 50 mM final concentration. After brief mixing, CHX was added to each tube at 200 μg/ml final concentration and then incubated for another 1 h at 37 °C before taking an aliquot for cell lysate preparation using the NaOH/TCA method (see preceding section). For the unbuffered control, an equal volume of ddH_2_O was added to the tube, and immediately after adding CHX, a 1 ml culture aliquot was mixed with NaOH to prepare cell lysate, which was then used as reference (T = 0 h) for all comparisons. The selection of pH buffers used was based on their buffering capacities: sodium acetate (pH = 4, 5), MES (pH = 6), and HEPES (pH = 7, 8).

### Glutamate analysis

Following manufacturer’s instructions, glutamate levels in cells or spent medium were analyzed in a microplate format using the Glutamate Assay Kit from Abcam (Fluorometric, LOQ ∼1 μM; Cat.#ab138883). **i. Total intracellular glutamate.** Cells (CFG404) from overnight YPD cultures were harvested, washed, and inoculated in YNB+CA without or with 2% glucose at OD ≍ 2. After 2 h of growth at 37 °C, cells were harvested, washed 3x with ice-cold ddH_2_O, and then resuspended in lysis buffer (25 mM Tris-HCl (pH=7.2), 150 mM NaCl, 5 mM MgCl_2_, 1% NP-40 (Igepal CA-630), and 5% glycerol) with 1 mM PMSF and cOmplete™ protease inhibitor. Cells were lysed by bead-beating using FastPrep^TM^ (60 s at 6.0 m/s) for 5 cycles with 3 min rest on ice between cycles. Lysates were clarified by centrifugation at 17000 *g* for 10 min at 4 °C, and then the supernatant was saved for glutamate analysis, protein content determination, and immunoblotting. Protein content was analyzed by the Bicinchoninic acid (BCA) assay (Sigma). Samples for immunoblotting were added with 2x SDS sample buffer and run. Glutamate and protein measurements were performed in a BioRad (Enspire) microplate reader. **iii. Extracellular glutamate.** For glutamate analysis in spent medium, cells from YPD cultures of wildtype (SC5314) and *gdh2-/-* (CFG279) strains were harvested, washed extensively (4x) with excess ddH_2_O to remove the residual YPD medium, and then resuspended in ddH_2_O. Cells were diluted at OD_600_ ≍ 2 in 20 ml of prewarmed YNB+PALAAG medium and then grown in a shaking incubator at 37 °C, and then after 3 h or 5 h, a 5 ml aliquot was taken for analysis. Briefly, the cultures were centrifuged at 4000 *g* for 5 min, and then a 3 ml aliquot of the supernatant was removed and filtered (0.2 μm). A 200 μl from this filtered supernatant and the sterile medium were neutralized with 10 μl of 1M HEPES, pH = 7.4 (50 mM final concentration) to convert all glutamic acids to glutamate. Except for the uninoculated medium, samples were diluted in 50 mM HEPES before subjecting to glutamate analysis. For practicality, samples collected on different days were stored at –80 °C until the run. Results presented were derived from 4-7 biological replicates performed in duplicates and run in a TECAN microplate reader in two batches (2 different days). **iii. Cytosolic glutamate.** To analyze cytosolic glutamate in cells treated with antimycin, we followed the cytosolic extraction protocol by Ohsumi et al. [20] with minor modifications. Briefly, cells (CFG441) harvested from overnight YPD cultures were washed and then diluted into 20 ml of YNB+CASG at OD ≍ 2. Cultures were grown for 2 h at 37 °C. Then, 5 ml aliquots of the culture were transferred into two 15-ml falcon tubes and immediately spiked with either antimycin A (1 μg/ml final concentration) or ethanol (vehicle). Cultures were placed in the 37 °C shaker for 30 min, and then the cells were harvested, washed 3x, and resuspended by brief vortexing in 1.5 ml of the extraction buffer [2.5 mM potassium phosphate (pH = 6), 0.6 M sorbitol, 10 mM glucose, and 0.2 mM CuCl_2_]. Cells were incubated at 30 °C for 10 min with gentle shaking (70 rpm) to permeabilize the cell membrane releasing the cytosolic amino acids. After incubation, a 1 ml aliquot was passed through a 0.45 μm filter connected to a syringe to collect the filtrate. An aliquot of the extract was neutralized by adding 1M HEPES (pH = 7.4) to 50 mM final concentration to convert glutamic acids to glutamate before analyzing for glutamate.

### Alpha-Ketoglutarate analysis

The cytosolic extracts obtained from antimycin-treated cells (see preceding section) were also analyzed for α-ketoglutarate using the Alpha Ketoglutarate (alpha KG) Assay Kit (Cat. # ab83431) following the manufacturer’s instruction. Briefly, aliquots of the extracts were first deproteinated by TCA and then later neutralized with KOH. Neutralized extracts were added with HEPES (to 50 mM final concentration) to ensure the extract pH was around 7.4 during analysis.

### Fluconazole susceptibility assay

Broth microdilution assay was performed in a microplate format according to the protocol by EUCAST method [57] with minor modifications that are limited to dissolving the fluconazole powder in 2% DMSO solution in ddH_2_O and cell density adjustment by OD (1 OD ≍ 3 × 10^7^ CFU/ml). The minimum inhibitory concentration (MIC) is the lowest drug concentration that gives rise to ≥50% growth inhibition of the drug-free control.

### *Ex vivo* human whole blood infection

Freshly extracted anonymized blood samples (4 ml) were purchased from Blodcentralen (Odenplan, Stockholm, Sweden), and all investigations were conducted according to the principles expressed in the Declaration of Helsinki. The well-established donation protocol for collecting human blood by the Karolinska University Laboratory for purposes other than medical treatment provided samples of peripheral blood collected from healthy volunteers with explicit consent for use in this study. According to legal requirements, the samples were labeled with donation numbers for traceability, but no names or other personal data was provided. Upon receipt in our laboratory, the samples are de-identified and can no longer be traced to an individual. This study does not require specific ethical approval.

For infection, fungal cells were first grown overnight in YPD and then refreshed the following day in a fresh medium. Exponentially growing cells were collected, washed twice in PBS, and then adjusted to 1 x 10^6^ CFU/ml. Around 1 x 10^5^ cells (10 μl) was added to 400 μl of whole blood (+EDTA), mixed briefly, and then incubated for 1 h in a shaking heat block (Eppendorf) set at 37 ℃ and 400 rpm. After incubation, tubes were vortexed vigorously, serially diluted in ddH_2_O (to lyse human cells), and plated on YPD agar. Plates were incubated at 30 ℃ for 2 days and colonies (CFU) counted. % Survival of fungal cells was calculated by comparing the CFU after the 1h incubation to the CFU of the inoculum.

### Transwell assay

*Lactobacillus crispatus* (LC100) vaginal isolate from a healthy female was recovered on MRS agar medium from glycerol stocks and incubated anaerobically for 48 h at 37 ℃ in anaerobic jar (GasPak™) with an Anaerogen sachet (Oxoid). For broth culture, isolated colonies were grown anaerobically in MRS broth for 48 h at 37 ℃. Cells were then collected by centrifugation 6000 *g* and then washed three times to remove the media. Cells were inoculated into CNY medium at ∼3 x10^7^ CFU/ml, and then a 500-μl aliquot (∼1.5 x10^7^ CFU/well) of the suspension was gently dispensed in the well of a 24-well plate containing a sterile insert (Millicell, 0.4 μm PCF, Cat. # PIHP01250; Merck Millipore, Ltd.) placed at the center of the well. For the processing of fungal cells, *cph1*Δ/Δ *efg1*Δ/Δ (CASJ041) or *cph1*Δ/Δ *efg1*Δ/Δ *gdh2*-/– (CFG352) cells were harvested from overnight YPD broth, washed twice in ddH_2_O, and then resuspended in CNY medium at OD ≍ 5. A 200 μl aliquot (OD ≍ 1; ∼1.6 x10^7^ CFU/well) was carefully pipetted into the center of the insert. Plates were then incubated in aerobic, microaerophilic, and anaerobic conditions for 48 h at 37 ℃ before photography and sampling the external culture for viable cell count via plating on MRS agar. All preparations were done in a Don Whitley anaerobic chamber (85% N_2_, 10% CO_2_, 5% H_2_) prior to incubation. Aerobic incubation was done in static, ambient conditions while microaerophilic (6.2-13.2% O_2,_ 2.5-9.5% CO_2_) and anaerobic (<1% O_2_, 9-13% CO_2_) conditions were met using an anaerobic jar with a Campygen sachet (Oxoid) and Anaerogen sachet (Oxoid), respectively.

### Statistical analysis

Data shown in this work were derived from at least 3 independent biological replicates and were analyzed using GraphPad Prism version 9. Specific statistical treatment applied and error bars used are described in the appropriate figure description and applied to the data obtained. Depending on the experiment, statistical treatments include unpaired student *t*-test or regular one-way analysis of variance (ANOVA) followed by Dunnett’s posthoc test. The following notations were used to describe statistical significance: \**p*<0.05, \*\**p*<0.01, \*\*\**p*<0.001, \*\*\*\**p*<0.0001, ns = not significant.

## Acknowledgments

The authors would like to thank the current and former members of the Per O. Ljungdahl, Claes Andréasson, and Sabrina Büttner laboratories (Stockholm University, Sweden) for their constructive comments throughout the course of this work. Gratitude is also extended to Kicki Ryman (Stockholm University, Sweden) and Sabrina Jenull (Medical University of Vienna, Austria) for fruitful discussions. We would also like to thank the following individuals for supplying strains: Valmik Vyas and Gerald R. Fink (MIT, USA), Karl Kuchler (Medical University of Vienna, Austria), Slavena Vylkova (Friedrich Schiller University, Germany), and Changbin Chen (Institute Pasteur of Shanghai, China). Lars Engstrand (Karolinska Institute, Sweden) is gratefully acknowledged for enabling the study of lactobacillus co-culture in his laboratory and for the productive discussions. **Funding:** This research was supported by: Swedish Research Council 2019-01547, 2022-01190 and Marie Curie – Initial Training Networks (ITN), ImResFun 606786 (POL).

## Author contributions

Conceptualization: FGSS, POL

Methodology: FGSS, EU, EF, VDV

Investigation: FGSS, EU, EF, VDV

Visualization: FGSS, EU, EF, VDV

Funding acquisition: POL

Supervision: POL, FGSS

Writing – original draft: FGSS

Writing – review & editing: FGSS, POL, VDV

## Data and materials availability

All data are available in the main text or the supplementary Table S1.

## Supplementary Materials

### Figures

Figure S1. Amino acid-dependent alkalization is dependent on Stp2 and Put3.

Figure S2. *ATO5* is dispensable for amino acid-dependent alkalization.

Figure S3. Viable cell count of *L. crispatus* grown for 48 h under the indicated oxygen tension.

### Tables

Table S1. Key resource table

---

**Fig. S1.**
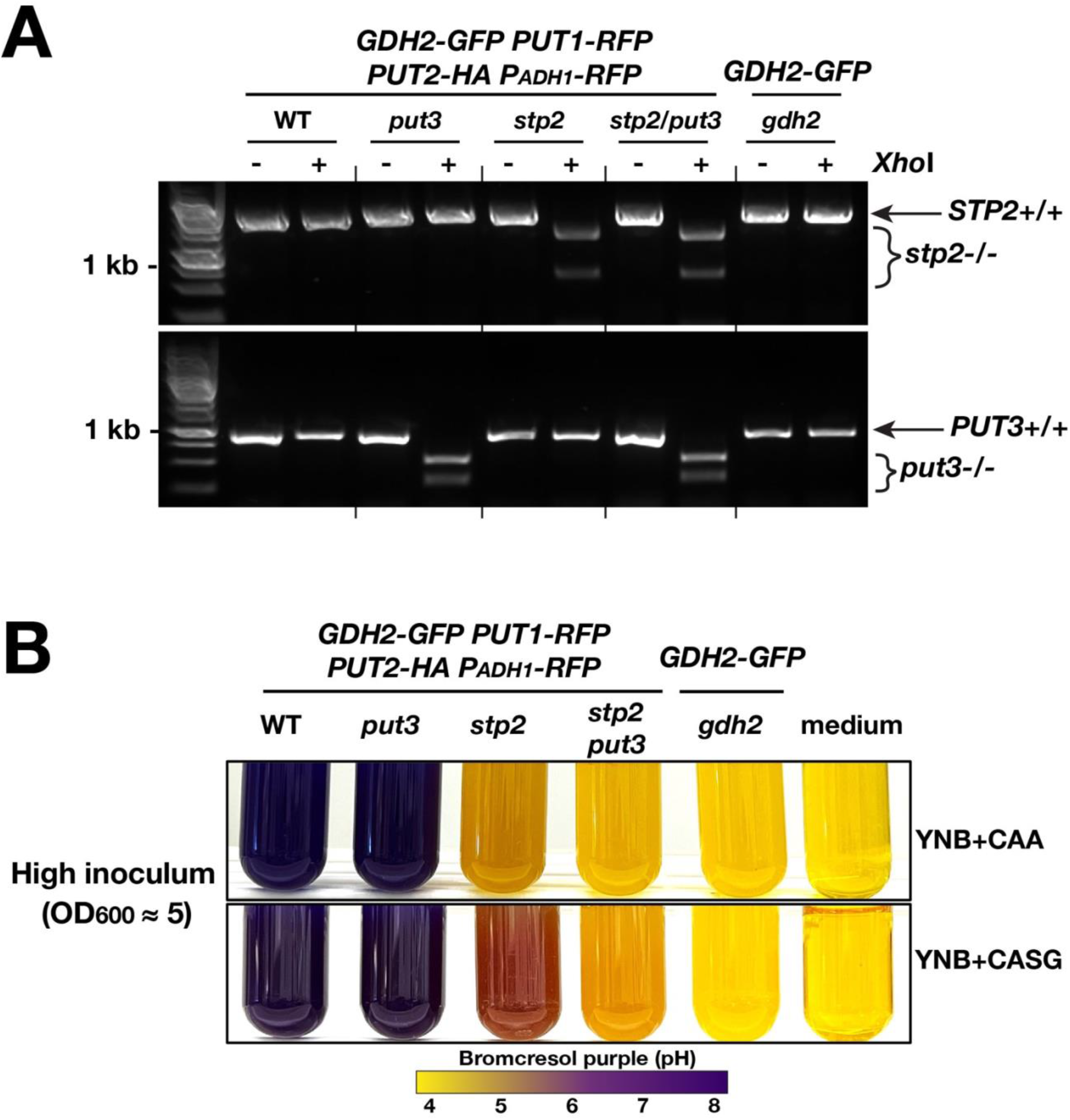
Amino acid-dependent alkalization is dependent on Stp2 and Put3. A) PCR and restriction digest verification of the *stp2-/-* and/or *put3-/-* strains showing the wildtype (+/+; arrow) and mutated genes digested by *Xho*I (–/-; bracket). Strains used: WT (CFG441), *put3* (CFG443), *stp2* (CFG665), *stp2 put3* (CFG671), and *gdh2* (CFG412). **B)** Cells of the same genotypes as (**A**) were grown in YNB+CASG at initial OD ≍ 5 and then photographed after 24 h.

**Fig. S2.**
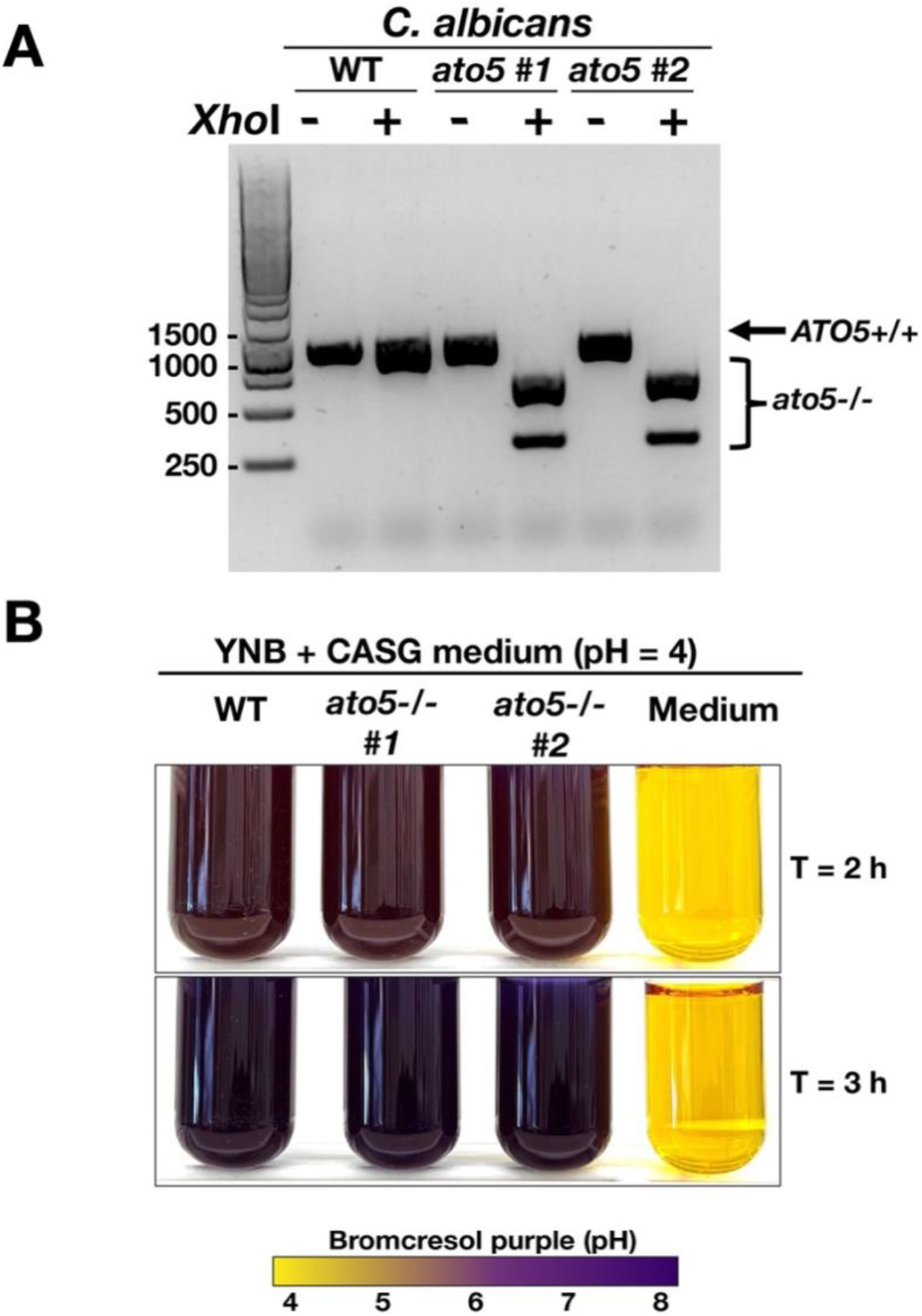
*ATO5* is dispensable for amino acid-dependent alkalization. A) PCR and restriction digest verification of the *ato5-/-* strains showing the wildtype (+/+; arrow) and mutated genes digested by *Xho*I (–/-; bracket). Strains used: WT (SC5314), *ato5-/– #1* (CFG691), *ato5-/– #2* (CFG694). **B)** Cells of the same genotypes as (A) were grown in YNB+CASG at initial OD ≍ 5 and then photographed at the indicated timepoints.

**Fig. S3.**
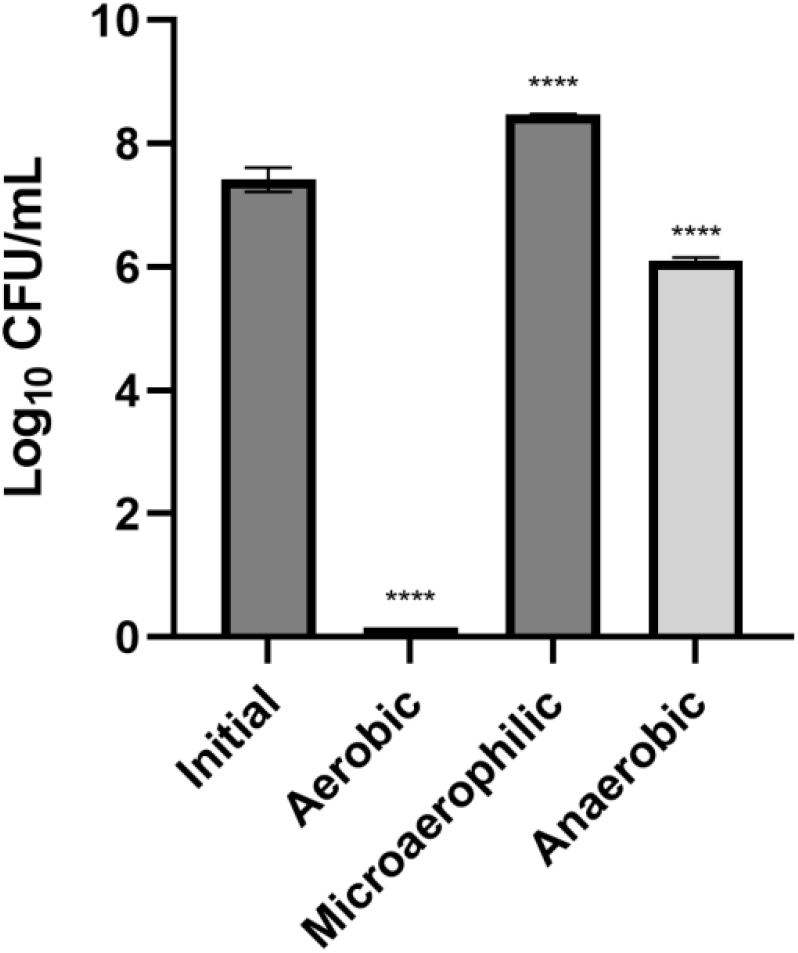
Viable cell count of *L. crispatus* grown for 48 h under the indicated oxygen tension. Data presented are mean of 4 biological replicates analyzed by one way ANOVA with Dunnett’s multiple comparison test. (****) represent an adjusted p-value of < 0.0001.

**Table S1.**
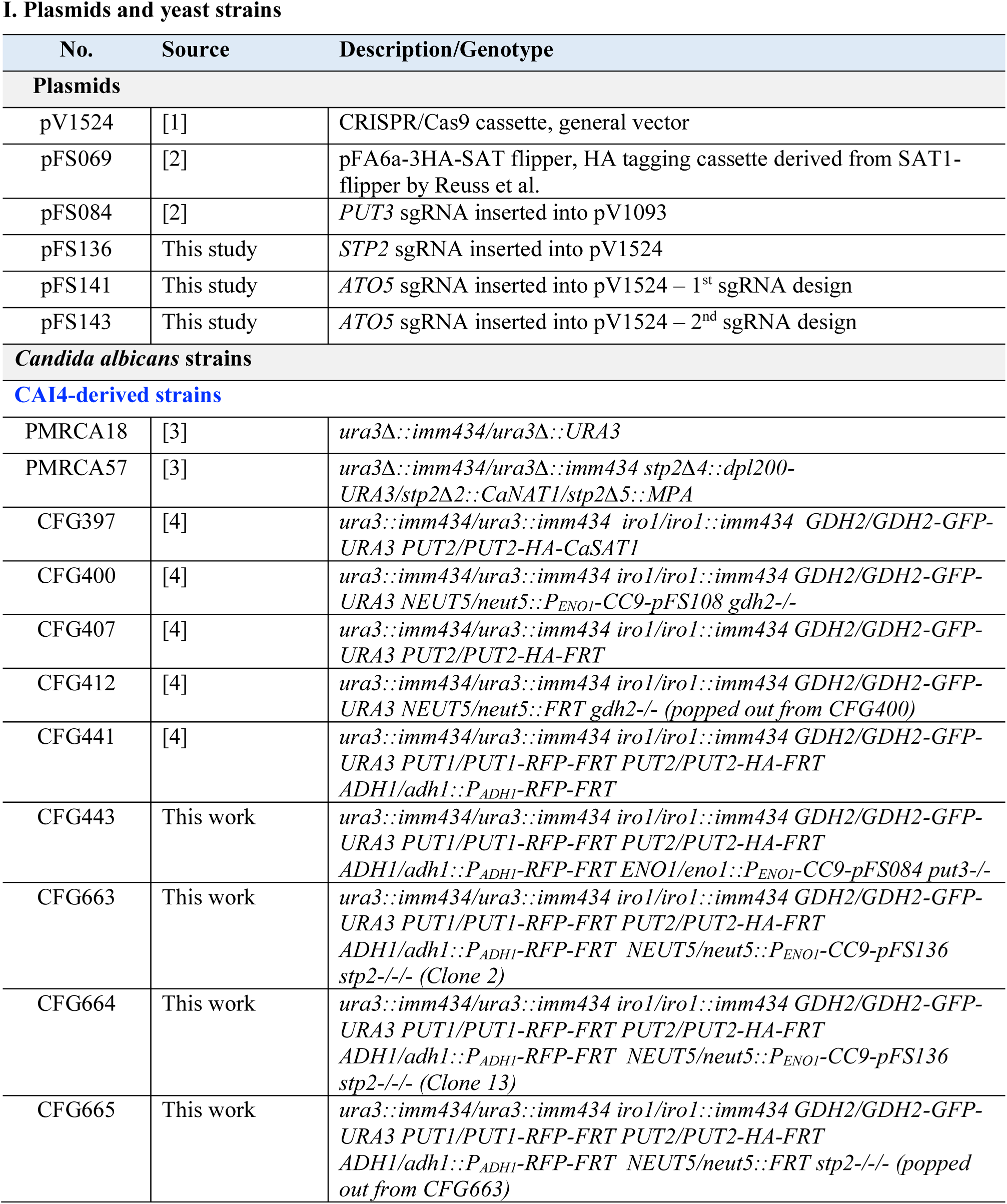
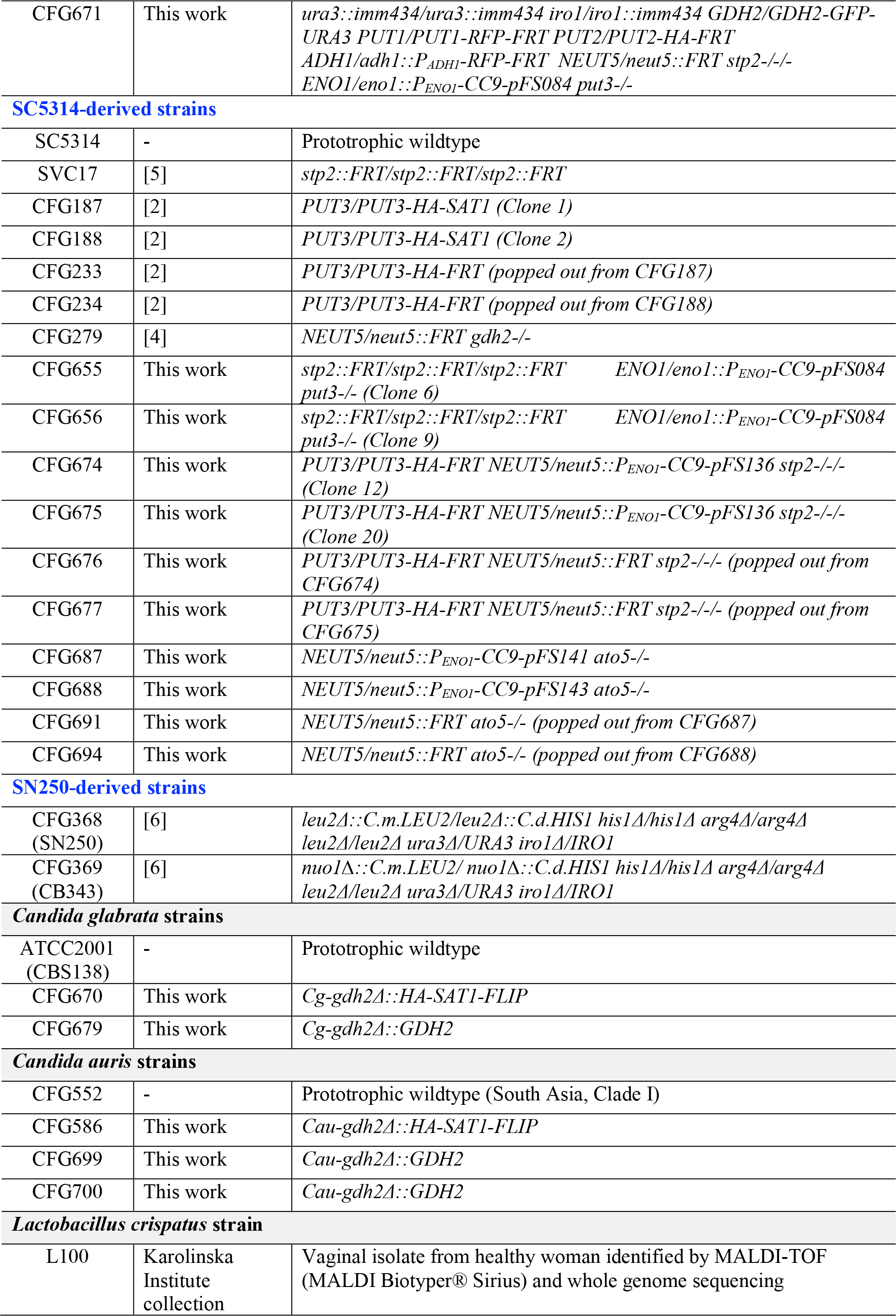
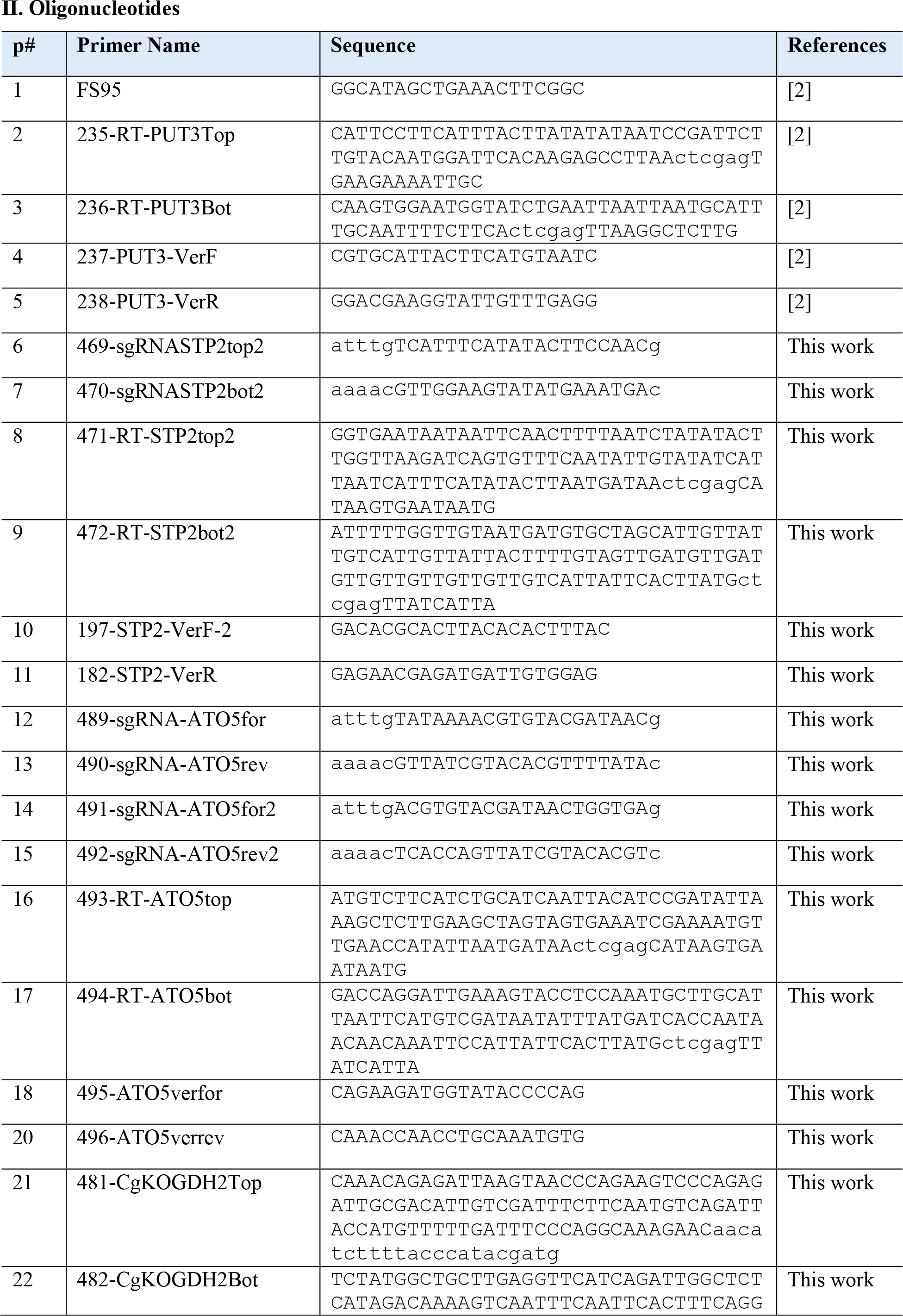
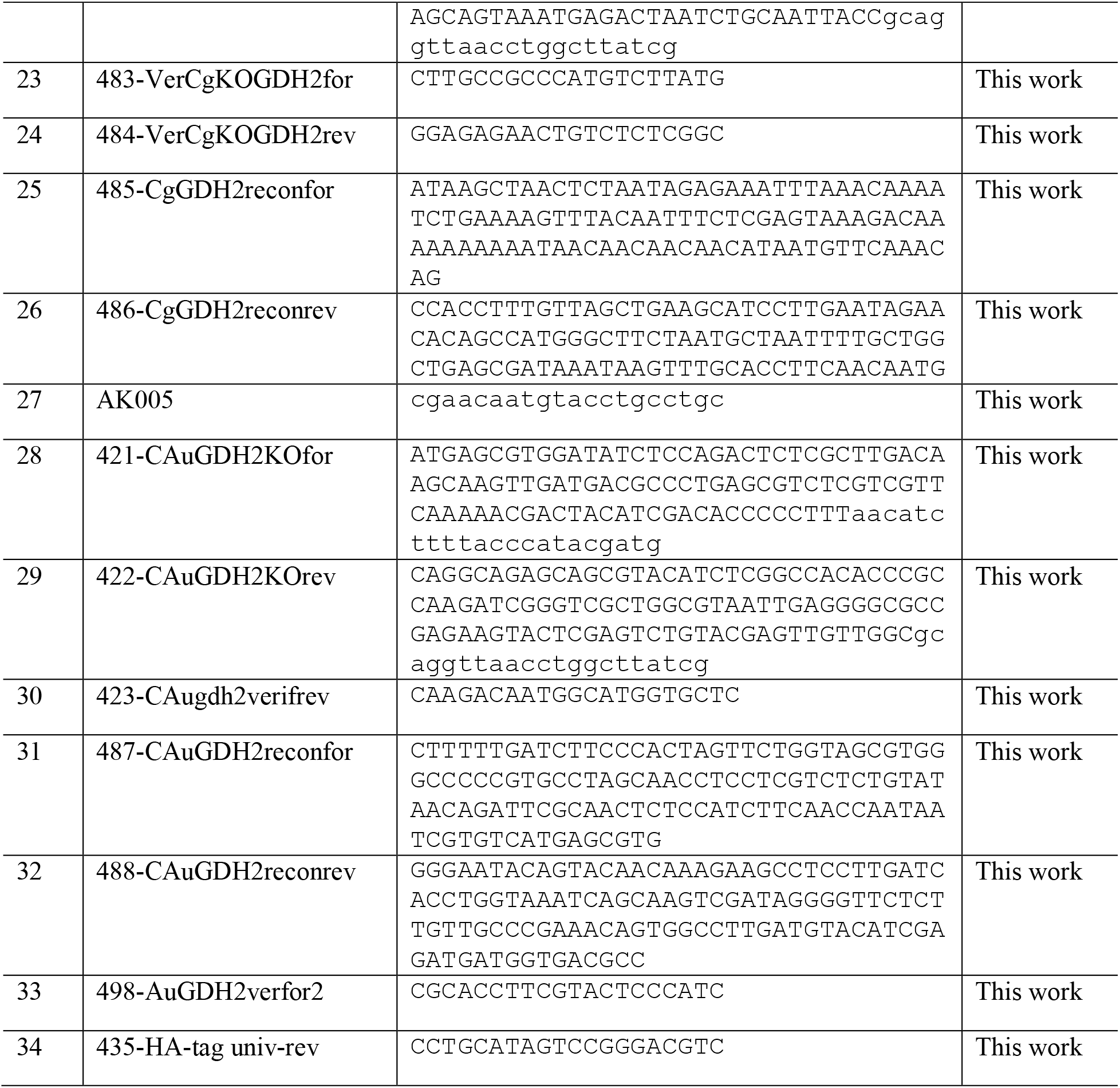
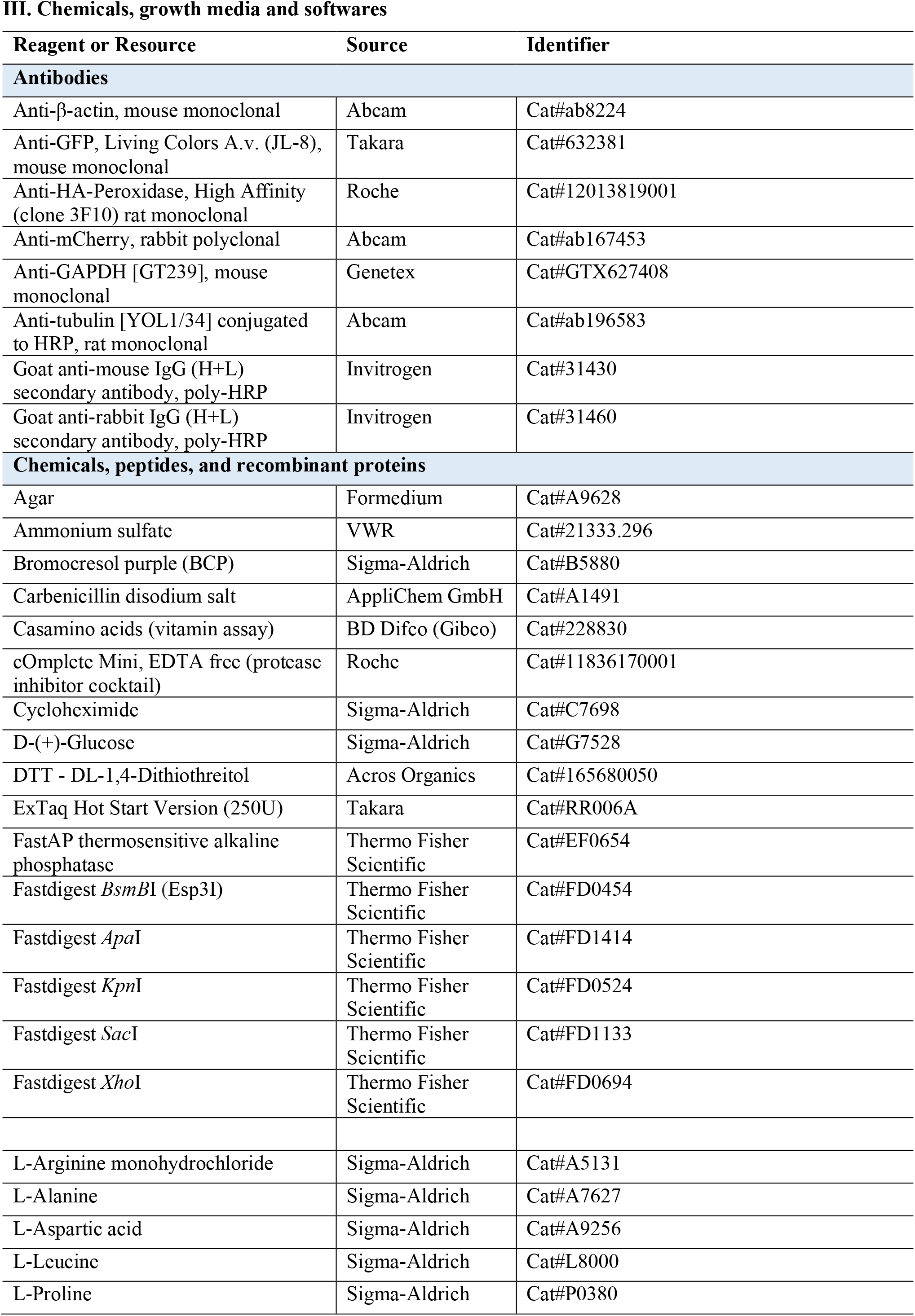
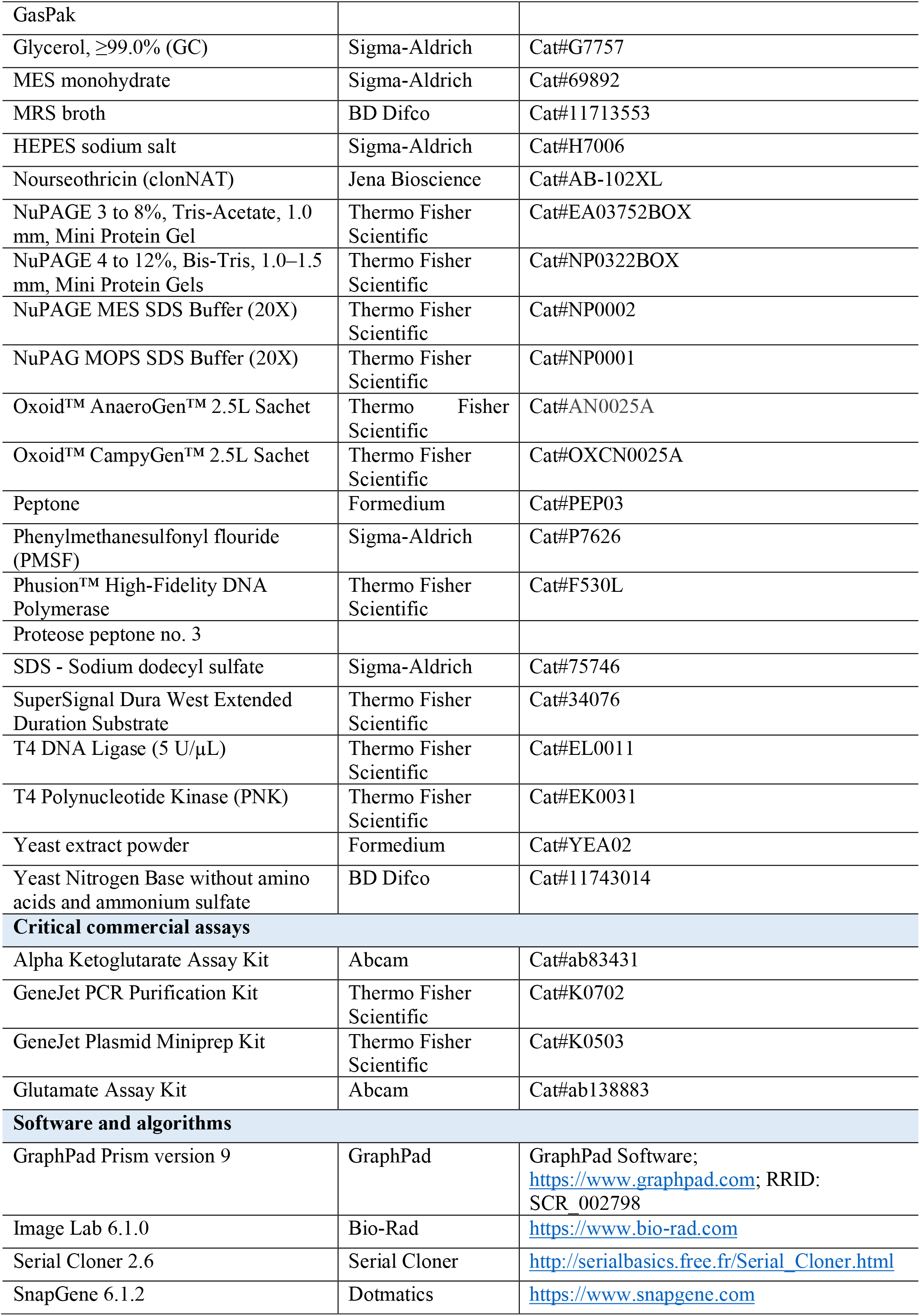
Key resource table. I. Plasmids and yeast strains
II. Oligonucleotides,
III. Chemicals, growth media and softwares

## Bibliography

1. Vylkova S. Environmental pH modulation by pathogenic fungi as a strategy to conquer the host. PLoS Pathog. 2017;13(2):e1006149. Epub 2017/02/24. doi: 10.1371/journal.ppat.1006149. PubMed PMID: 28231317; PubMed Central PMCID: PMCPMC5322887.

2. Vylkova S, Carman AJ, Danhof HA, Collette JR, Zhou H, Lorenz MC. The fungal pathogen Candida albicans autoinduces hyphal morphogenesis by raising extracellular pH. mBio. 2011;2(3):e00055–11. doi: 10.1128/mBio.00055-11. PubMed PMID: 21586647; PubMed Central PMCID: PMC3101780.

3. WHO fungal priority pathogens list to guide research, development and public health action. Geneva: World Health Organization. 2022.

4. Fisher MC, Denning DW. The WHO fungal priority pathogens list as a game-changer. Nat Rev Microbiol. 2023;21(4):211–2. doi: 10.1038/s41579-023-00861-x. PubMed PMID: 36747091; PubMed Central PMCID: PMCPMC9901396.

5. Silao FGS, Ward M, Ryman K, Wallstrom A, Brindefalk B, Udekwu K, et al. Mitochondrial proline catabolism activates Ras1/cAMP/PKA-induced filamentation in Candida albicans. PLoS Genet. 2019;15(2):e1007976. Epub 2019/02/12. doi: 10.1371/journal.pgen.1007976. PubMed PMID: 30742618.

6. Silao FGS, Ryman K, Jiang T, Ward M, Hansmann N, Molenaar C, et al. Glutamate dehydrogenase (Gdh2)-dependent alkalization is dispensable for escape from macrophages and virulence of Candida albicans. PLoS Pathog. 2020;16(9):e1008328. Epub 2020/09/17. doi: 10.1371/journal.ppat.1008328. PubMed PMID: 32936835; PubMed Central PMCID: PMCPMC7521896.

7. Han TL, Cannon RD, Gallo SM, Villas-Boas SG. A metabolomic study of the effect of Candida albicans glutamate dehydrogenase deletion on growth and morphogenesis. NPJ Biofilms Microbiomes. 2019;5:13. Epub 2019/04/18. doi: 10.1038/s41522-019-0086-5. PubMed PMID: 30992998; PubMed Central PMCID: PMCPMC6453907 non-financial relationships that could be construed as a potential conflict of interest.

8. Vylkova S, Lorenz MC. Modulation of phagosomal pH by Candida albicans promotes hyphal morphogenesis and requires Stp2p, a regulator of amino acid transport. PLoS Pathog. 2014;10(3):e1003995. doi: 10.1371/journal.ppat.1003995. PubMed PMID: 24626429; PubMed Central PMCID: PMC3953444.

9. Danhof HA, Vylkova S, Vesely EM, Ford AE, Gonzalez-Garay M, Lorenz MC. Robust Extracellular pH Modulation by Candida albicans during Growth in Carboxylic Acids. mBio. 2016;7(6). Epub 2016/12/10. doi: 10.1128/mBio.01646-16. PubMed PMID: 27935835; PubMed Central PMCID: PMCPMC5111404.

10. Miramon P, Lorenz MC. The SPS amino acid sensor mediates nutrient acquisition and immune evasion in Candida albicans. Cell Microbiol. 2016;18(11):1611–24. Epub 2016/10/26. doi: 10.1111/cmi.12600. PubMed PMID: 27060451; PubMed Central PMCID: PMCPMC5501722.

11. Miramon P, Pountain AW, van Hoof A, Lorenz MC. The Paralogous Transcription Factors Stp1 and Stp2 of Candida albicans Have Distinct Functions in Nutrient Acquisition and Host Interaction. Infect Immun. 2020;88(5). Epub 20200420. doi: 10.1128/IAI.00763-19. PubMed PMID: 32094252; PubMed Central PMCID: PMCPMC7171245.

12. Hollomon JM, Liu Z, Rusin SF, Jenkins NP, Smith AK, Koeppen K, et al. The Candida albicans Cdk8-dependent phosphoproteome reveals repression of hyphal growth through a Flo8-dependent pathway. PLoS Genet. 2022;18(1):e1009622. Epub 20220104. doi: 10.1371/journal.pgen.1009622. PubMed PMID: 34982775; PubMed Central PMCID: PMCPMC8769334.

13. Todd OA, Noverr MC, Peters BM. Candida albicans Impacts Staphylococcus aureus Alpha-Toxin Production via Extracellular Alkalinization. mSphere. 2019;4(6). Epub 20191113. doi: 10.1128/mSphere.00780-19. PubMed PMID: 31722996; PubMed Central PMCID: PMCPMC6854045.

14. Martinez P, Ljungdahl PO. Divergence of Stp1 and Stp2 transcription factors in Candida albicans places virulence factors required for proper nutrient acquisition under amino acid control. Mol Cell Biol. 2005;25(21):9435–46. doi: 10.1128/MCB.25.21.9435-9446.2005. PubMed PMID: 16227594; PubMed Central PMCID: PMC1265835.

15. Amorim-Vaz S, Delarze E, Ischer F, Sanglard D, Coste AT. Examining the virulence of Candida albicans transcription factor mutants using Galleria mellonella and mouse infection models. Front Microbiol. 2015;6:367. Epub 2015/05/23. doi: 10.3389/fmicb.2015.00367. PubMed PMID: 25999923; PubMed Central PMCID: PMCPMC4419840.

16. Danhof HA, Lorenz MC. The Candida albicans ATO Gene Family Promotes Neutralization of the Macrophage Phagolysosome. Infect Immun. 2015;83(11):4416–26. doi: 10.1128/IAI.00984-15. PubMed PMID: 26351284; PubMed Central PMCID: PMCPMC4598414.

17. Silao FGS, Jiang T, Bereczky-Veress B, Kühbacher A, Ryman K, Uwamohoro N, et al. Proline catabolism is key to facilitating Candida albicans pathogenicity. bioRxiv. 2023:2023.01.17.524449. doi: 10.1101/2023.01.17.524449.

18. Hebert EM, Mamone G, Picariello G, Raya RR, Savoy G, Ferranti P, et al. Characterization of the pattern of alphas1– and beta-casein breakdown and release of a bioactive peptide by a cell envelope proteinase from Lactobacillus delbrueckii subsp. lactis CRL 581. Appl Environ Microbiol. 2008;74(12):3682–9. Epub 20080418. doi: 10.1128/AEM.00247-08. PubMed PMID: 18424544; PubMed Central PMCID: PMCPMC2446539.

19. Silao FGS, Ljungdahl PO. Amino Acid Sensing and Assimilation by the Fungal Pathogen Candida albicans in the Human Host. Pathogens. 2021;11(1). Epub 20211222. doi: 10.3390/pathogens11010005. PubMed PMID: 35055954; PubMed Central PMCID: PMCPMC8781990.

20. Ohsumi Y, Kitamoto K, Anraku Y. Changes induced in the permeability barrier of the yeast plasma membrane by cupric ion. J Bacteriol. 1988;170(6):2676–82. PubMed PMID: 3286617; PubMed Central PMCID: PMC211187.

21. Huang X, Chen X, He Y, Yu X, Li S, Gao N, et al. Mitochondrial complex I bridges a connection between regulation of carbon flexibility and gastrointestinal commensalism in the human fungal pathogen Candida albicans. PLoS Pathog. 2017;13(6):e1006414. Epub 20170601. doi: 10.1371/journal.ppat.1006414. PubMed PMID: 28570675; PubMed Central PMCID: PMCPMC5469625.

22. She X, Khamooshi K, Gao Y, Shen Y, Lv Y, Calderone R, et al. Fungal-specific subunits of the Candida albicans mitochondrial complex I drive diverse cell functions including cell wall synthesis. Cell Microbiol. 2015;17(9):1350–64. Epub 2015/03/25. doi: 10.1111/cmi.12438. PubMed PMID: 25801605; PubMed Central PMCID: PMCPMC4677794.

23. Velasco I, Tenreiro S, Calderon IL, Andre B. Saccharomyces cerevisiae Aqr1 Is an Internal-Membrane Transporter Involved in Excretion of Amino Acids. Eukaryotic Cell. 2004;3(6):1492–503. doi: 10.1128/ec.3.6.1492-1503.2004.

24. Bottcher B, Driesch D, Kruger T, Garbe E, Gerwien F, Kniemeyer O, et al. Impaired amino acid uptake leads to global metabolic imbalance of Candida albicans biofilms. NPJ Biofilms Microbiomes. 2022;8(1):78. Epub 20221013. doi: 10.1038/s41522-022-00341-9. PubMed PMID: 36224215; PubMed Central PMCID: PMCPMC9556537.

25. Teixeira MC, Viana R, Palma M, Oliveira J, Galocha M, Mota MN, et al. YEASTRACT+: a portal for the exploitation of global transcription regulation and metabolic model data in yeast biotechnology and pathogenesis. Nucleic Acids Res. 2023;51(D1):D785–D91. doi: 10.1093/nar/gkac1041. PubMed PMID: 36350610; PubMed Central PMCID: PMCPMC9825512.

26. Tebung WA, Omran RP, Fulton DL, Morschhauser J, Whiteway M. Put3 Positively Regulates Proline Utilization in Candida albicans. mSphere. 2017;2(6). Epub 2017/12/16. doi: 10.1128/mSphere.00354-17. PubMed PMID: 29242833; PubMed Central PMCID: PMCPMC5729217.

27. de Boer CG, Hughes TR. YeTFaSCo: a database of evaluated yeast transcription factor sequence specificities. Nucleic Acids Res. 2012;40(Database issue):D169–79. Epub 20111118. doi: 10.1093/nar/gkr993. PubMed PMID: 22102575; PubMed Central PMCID: PMCPMC3245003.

28. Swaminathan K, Flynn P, Reece RJ, Marmorstein R. Crystal structure of a PUT3-DNA complex reveals a novel mechanism for DNA recognition by a protein containing a Zn2Cys6 binuclear cluster. Nat Struct Biol. 1997;4(9):751–9. doi: 10.1038/nsb0997-751. PubMed PMID: 9303004.

29. de Boer M, Nielsen PS, Bebelman JP, Heerikhuizen H, Andersen HA, Planta RJ. Stp1p, Stp2p and Abf1p are involved in regulation of expression of the amino acid transporter gene BAP3 of Saccharomyces cerevisiae. Nucleic Acids Res. 2000;28(4):974–81. doi: 10.1093/nar/28.4.974. PubMed PMID: 10648791; PubMed Central PMCID: PMCPMC102570.

30. Yang TH, Wu WS. Identifying biologically interpretable transcription factor knockout targets by jointly analyzing the transcription factor knockout microarray and the ChIP-chip data. BMC Syst Biol. 2012;6:102. Epub 20120816. doi: 10.1186/1752-0509-6-102. PubMed PMID: 22898448; PubMed Central PMCID: PMCPMC3465233.

31. Lagree K, Woolford CA, Huang MY, May G, McManus CJ, Solis NV, et al. Roles of Candida albicans Mig1 and Mig2 in glucose repression, pathogenicity traits, and SNF1 essentiality. PLoS Genet. 2020;16(1):e1008582. Epub 2020/01/22. doi: 10.1371/journal.pgen.1008582. PubMed PMID: 31961865; PubMed Central PMCID: PMCPMC6994163.

32. Smith DFQ, Mudrak NJ, Zamith-Miranda D, Honorato L, Nimrichter L, Chrissian C, et al. Melanization of Candida auris Is Associated with Alteration of Extracellular pH. J Fungi (Basel). 2022;8(10). Epub 20221011. doi: 10.3390/jof8101068. PubMed PMID: 36294632; PubMed Central PMCID: PMCPMC9604884.

33. Kasper L, Seider K, Gerwien F, Allert S, Brunke S, Schwarzmuller T, et al. Identification of Candida glabrata genes involved in pH modulation and modification of the phagosomal environment in macrophages. PloS one. 2014;9(5):e96015. Epub 2014/05/03. doi: 10.1371/journal.pone.0096015. PubMed PMID: 24789333; PubMed Central PMCID: PMCPMC4006850.

34. Reuss O, Vik A, Kolter R, Morschhauser J. The SAT1 flipper, an optimized tool for gene disruption in Candida albicans. Gene. 2004;341:119–27. doi: 10.1016/j.gene.2004.06.021. PubMed PMID: 15474295.

35. Fukasawa Y, Tsuji J, Fu SC, Tomii K, Horton P, Imai K. MitoFates: improved prediction of mitochondrial targeting sequences and their cleavage sites. Mol Cell Proteomics. 2015;14(4):1113–26. Epub 2015/02/12. doi: 10.1074/mcp.M114.043083. PubMed PMID: 25670805; PubMed Central PMCID: PMCPMC4390256.

36. Kammer P, McNamara S, Wolf T, Conrad T, Allert S, Gerwien F, et al. Survival Strategies of Pathogenic Candida Species in Human Blood Show Independent and Specific Adaptations. mBio. 2020;11(5). Epub 20201006. doi: 10.1128/mBio.02435-20. PubMed PMID: 33024045; PubMed Central PMCID: PMCPMC7542370.

37. McCarthy MW, Walsh TJ. Amino Acid Metabolism and Transport Mechanisms as Potential Antifungal Targets. Int J Mol Sci. 2018;19(3). Epub 20180319. doi: 10.3390/ijms19030909. PubMed PMID: 29562716; PubMed Central PMCID: PMCPMC5877770.

38. Wang S, Wang Q, Yang E, Yan L, Li T, Zhuang H. Antimicrobial Compounds Produced by Vaginal Lactobacillus crispatus Are Able to Strongly Inhibit Candida albicans Growth, Hyphal Formation and Regulate Virulence-related Gene Expressions. Front Microbiol. 2017;8:564. Epub 20170404. doi: 10.3389/fmicb.2017.00564. PubMed PMID: 28421058; PubMed Central PMCID: PMCPMC5378977.

39. Sobel JD, Faro S, Force RW, Foxman B, Ledger WJ, Nyirjesy PR, et al. Vulvovaginal candidiasis: epidemiologic, diagnostic, and therapeutic considerations. Am J Obstet Gynecol. 1998;178(2):203–11. doi: 10.1016/s0002-9378(98)80001-x. PubMed PMID: 9500475.

40. Antonio MA, Hawes SE, Hillier SL. The identification of vaginal Lactobacillus species and the demographic and microbiologic characteristics of women colonized by these species. The Journal of infectious diseases. 1999;180(6):1950–6. doi: 10.1086/315109. PubMed PMID: 10558952.

41. Ehrstrom S, Yu A, Rylander E. Glucose in vaginal secretions before and after oral glucose tolerance testing in women with and without recurrent vulvovaginal candidiasis. Obstet Gynecol. 2006;108(6):1432–7. doi: 10.1097/01.AOG.0000246800.38892.fc. PubMed PMID: 17138777.

42. Miller EA, Beasley DE, Dunn RR, Archie EA. Lactobacilli Dominance and Vaginal pH: Why Is the Human Vaginal Microbiome Unique? Front Microbiol. 2016;7:1936. Epub 20161208. doi: 10.3389/fmicb.2016.01936. PubMed PMID: 28008325; PubMed Central PMCID: PMCPMC5143676.

43. Fernandes TR, Segorbe D, Prusky D, Di Pietro A. How alkalinization drives fungal pathogenicity. PLoS Pathog. 2017;13(11):e1006621. Epub 2017/11/10. doi: 10.1371/journal.ppat.1006621. PubMed PMID: 29121119; PubMed Central PMCID: PMCPMC5679519.

44. Miramon P, Lorenz MC. A feast for Candida: Metabolic plasticity confers an edge for virulence. PLoS Pathog. 2017;13(2):e1006144. doi: 10.1371/journal.ppat.1006144. PubMed PMID: 28182769.

45. Navarathna DH, Harris SD, Roberts DD, Nickerson KW. Evolutionary aspects of urea utilization by fungi. FEMS Yeast Res. 2010;10(2):209–13. Epub 20100121. doi: 10.1111/j.1567-1364.2010.00602.x. PubMed PMID: 20100286; PubMed Central PMCID: PMCPMC2822880.

46. Cueto-Rojas HF, Milne N, van Helmond W, Pieterse MM, van Maris AJA, Daran JM, et al. Membrane potential independent transport of NH3 in the absence of ammonium permeases in Saccharomyces cerevisiae. BMC Syst Biol. 2017;11(1):49. Epub 2017/04/18. doi: 10.1186/s12918-016-0381-1. PubMed PMID: 28412970; PubMed Central PMCID: PMCPMC5392931.

47. Labotka RJ, Lundberg P, Kuchel PW. Ammonia permeability of erythrocyte membrane studied by 14N and 15N saturation transfer NMR spectroscopy. Am J Physiol. 1995;268(3 Pt 1):C686–99. Epub 1995/03/01. doi: 10.1152/ajpcell.1995.268.3.C686. PubMed PMID: 7900775.

48. Antonenko YN, Pohl P, Denisov GA. Permeation of ammonia across bilayer lipid membranes studied by ammonium ion selective microelectrodes. Biophys J. 1997;72(5):2187–95. Epub 1997/05/01. doi: 10.1016/S0006-3495(97)78862-3. PubMed PMID: 9129821; PubMed Central PMCID: PMCPMC1184413.

49. Morales DK, Grahl N, Okegbe C, Dietrich LE, Jacobs NJ, Hogan DA. Control of Candida albicans metabolism and biofilm formation by Pseudomonas aeruginosa phenazines. mBio. 2013;4(1):e00526–12. doi: 10.1128/mBio.00526-12. PubMed PMID: 23362320; PubMed Central PMCID: PMCPMC3560528.

50. Zhang Y, Pelechano V. High-throughput 5’P sequencing enables the study of degradation-associated ribosome stalls. Cell Rep Methods. 2021;1(1):100001. Epub 20210402. doi: 10.1016/j.crmeth.2021.100001. PubMed PMID: 35474692; PubMed Central PMCID: PMCPMC9017187.

51. Eastment MC, Balkus JE, Richardson BA, Srinivasan S, Kimani J, Anzala O, et al. Association Between Vaginal Bacterial Microbiota and Vaginal Yeast Colonization. The Journal of infectious diseases. 2021;223(5):914–23. doi: 10.1093/infdis/jiaa459. PubMed PMID: 32726445; PubMed Central PMCID: PMCPMC7938175.

52. van de Wijgert JH, Borgdorff H, Verhelst R, Crucitti T, Francis S, Verstraelen H, et al. The vaginal microbiota: what have we learned after a decade of molecular characterization? PloS one. 2014;9(8):e105998. Epub 2014/08/26. doi: 10.1371/journal.pone.0105998. PubMed PMID: 25148517; PubMed Central PMCID: PMCPMC4141851.

53. Vazquez-Munoz R, Dongari-Bagtzoglou A. Anticandidal Activities by Lactobacillus Species: An Update on Mechanisms of Action. Front Oral Health. 2021;2:689382. Epub 20210716. doi: 10.3389/froh.2021.689382. PubMed PMID: 35048033; PubMed Central PMCID: PMCPMC8757823.

54. Lin YP, Chen WC, Cheng CM, Shen CJ. Vaginal pH Value for Clinical Diagnosis and Treatment of Common Vaginitis. Diagnostics (Basel). 2021;11(11). Epub 20211027. doi: 10.3390/diagnostics11111996. PubMed PMID: 34829343; PubMed Central PMCID: PMCPMC8618584.

55. Wagner G, Levin R. Oxygen tension of the vaginal surface during sexual stimulation in the human. Fertil Steril. 1978;30(1):50–3. doi: 10.1016/s0015-0282(16)43395-9. PubMed PMID: 581075.

56. Malcher M, Schladebeck S, Mosch HU. The Yak1 protein kinase lies at the center of a regulatory cascade affecting adhesive growth and stress resistance in Saccharomyces cerevisiae. Genetics. 2011;187(3):717–30. Epub 20101213. doi: 10.1534/genetics.110.125708. PubMed PMID: 21149646; PubMed Central PMCID: PMCPMC3063667.

57. Subcommittee on Antifungal Susceptibility Testing of the EECfAST. EUCAST definitive document EDef 7.1: method for the determination of broth dilution MICs of antifungal agents for fermentative yeasts. Clin Microbiol Infect. 2008;14(4):398–405. Epub 20080111. doi: 10.1111/j.1469-0691.2007.01935.x. PubMed PMID: 18190574.

58. Erickson RJ. An Evaluation of Mathematical Models for the Effects of pH and Temperature on Ammonia Toxicity to Aquatic Organisms. Water Research. 1985;19(8):1047–58.

## References

1. Vyas VK, Bushkin GG, Bernstein DA, Getz MA, Sewastianik M, Barrasa MI, et al. New CRISPR Mutagenesis Strategies Reveal Variation in Repair Mechanisms among Fungi. mSphere. 2018;3(2). Epub 2018/04/27. doi: 10.1128/mSphere.00154-18. PubMed PMID: 29695624; PubMed Central PMCID: PMCPMC5917429.

2. Silao FGS, Ward M, Ryman K, Wallstrom A, Brindefalk B, Udekwu K, et al. Mitochondrial proline catabolism activates Ras1/cAMP/PKA-induced filamentation in Candida albicans. PLoS Genet. 2019;15(2):e1007976. Epub 2019/02/12. doi: 10.1371/journal.pgen.1007976. PubMed PMID: 30742618.

3. Martinez P, Ljungdahl PO. Divergence of Stp1 and Stp2 transcription factors in Candida albicans places virulence factors required for proper nutrient acquisition under amino acid control. Mol Cell Biol. 2005;25(21):9435–46. doi: 10.1128/MCB.25.21.9435-9446.2005. PubMed PMID: 16227594; PubMed Central PMCID: PMC1265835.

4. Silao FGS, Ryman K, Jiang T, Ward M, Hansmann N, Molenaar C, et al. Glutamate dehydrogenase (Gdh2)-dependent alkalization is dispensable for escape from macrophages and virulence of Candida albicans. PLoS Pathog. 2020;16(9):e1008328. Epub 2020/09/17. doi: 10.1371/journal.ppat.1008328. PubMed PMID: 32936835; PubMed Central PMCID: PMCPMC7521896.

5. Vylkova S, Lorenz MC. Modulation of phagosomal pH by Candida albicans promotes hyphal morphogenesis and requires Stp2p, a regulator of amino acid transport. PLoS Pathog. 2014;10(3):e1003995. doi: 10.1371/journal.ppat.1003995. PubMed PMID: 24626429; PubMed Central PMCID: PMC3953444.

6. Huang X, Chen X, He Y, Yu X, Li S, Gao N, et al. Mitochondrial complex I bridges a connection between regulation of carbon flexibility and gastrointestinal commensalism in the human fungal pathogen Candida albicans. PLoS Pathog. 2017;13(6):e1006414. Epub 20170601. doi: 10.1371/journal.ppat.1006414. PubMed PMID: 28570675; PubMed Central PMCID: PMCPMC5469625.

